# Systematic evaluation of blood contamination in nanoparticle-based plasma proteomics

**DOI:** 10.1101/2025.04.26.650757

**Authors:** Huanhuan Gao, Yuecheng Zhan, Yuanqi Liu, Zhiyi Zhu, Yuxiu Zheng, Liqin Qian, Zhangzhi Xue, Honghan Cheng, Zongxiang Nie, Weigang Ge, Senlin Ruan, Jiaxu Liu, Jikai Zhang, Yingying Sun, Lei Zhou, Dongyue Xun, Yingrui Wang, Heyun Xu, Huiwen Miao, Yi Zhu, Tiannan Guo

**Affiliations:** Affiliated Hangzhou First People’s Hospital, State Key Laboratory of Medical Proteomics, School of Medicine, Westlake University, Hangzhou, Zhejiang Province, China; Westlake Center for Intelligent Proteomics, Westlake Laboratory of Life Sciences and Biomedicine, Hangzhou, Zhejiang Province, China; Research Center for Industries of the Future, School of Life Sciences, Westlake University, Hangzhou, Zhejiang Province, China; Westlake Omics (Hangzhou) Biotechnology Co., Ltd., Hangzhou, China; The First Affiliated Hospital, Zhejiang University School of Medicine.; Department of Clinical Laboratory, Affiliated Hangzhou First People’s Hospital, Hangzhou, Zhejiang Province, China; State Key Laboratory of Fine Chemicals, Frontier Science Center for Smart Materials, School of Chemical Engineering, Dalian University of Technology, Dalian, China; College of Chemistry, Nankai University, Tianjin, China

**Keywords:** nanoparticle, silica, plasma proteomics, platelet contamination, erythrocyte contamination, coagulation contamination, lung cancer, biomarkers, Astral, EDTA tube, sodium citrate tube, lithium heparin tube

## Abstract

Circulating blood proteomics enables minimally invasive biomarker discovery. Nanoparticle (NP)-based circulating plasma proteomics studies have reported varying number of proteins, ranging from ca 2000 to 7000 but it’s not clear whether higher protein number is more informative. Here, we first develop OmniProt – a silica-NP workflow for plasma proteomics which is optimized through systematic evaluation of NP types and protein corona formation parameters. Next, we present an Astral spectral library for 10,109 protein groups expressed in human plasma. Using Astral with a throughput of 60 sample-per-day, OmniProt identifies ca 3000 to 6000 protein groups from human plasma. Notably, we found that platelet/erythrocyte/coagulation-related contamination artificially elevates protein identifications and compromises quantification accuracy for circulating plasma proteins in NP-enriched samples. Through controlled contamination experiments, we identified and validated protein biomarkers for contaminations of platelet, erythrocyte, and coagulation, respectively, in NP-based plasma proteomics experiments. We further developed an open-access software Baize for accessing plasma proteomics contamination. Finally, we validated our OmniProt workflow and Baize software in a clinical cohort of 193 patients with CT-indistinct benign nodules or early lung cancers. On average, we identified ca 4400 protein groups for each plasma sample without significant contamination. Five samples were flagged out as contaminated. In conclusion, this study reveals that contamination of plasma samples significantly alters the protein identification and quantification in NP-based plasma proteomics experiments, and presents an open-access software Baize to evaluate plasma proteome contamination.

## Introduction

Circulating blood proteomics has emerged as a pivotal research field for discovering disease-related biomarkers, due to sampling accessibility and the rich molecular insights derived from blood constituents^1–3^. The largest proteome study of about 600,000 circulating blood samples collected in UK Biobank has been initiated^4^ and supported by multiple pharmaceutical companies, as encouraged by the success of a pilot study of about 50,000 blood proteomes measured by Proximity Extension Assay^5–7^. The SomaLogic slow off-rate aptamer (SOMAmers)^8^ have also been used for thousands of samples^9, 10^. Unlike affinity-based technologies requiring a predefined protein target panel, mass spectrometry (MS)-based proteomics enables unbiased discovery without *a priori* protein selection^11^. But MS-based analysis suffers from limited proteome coverage due to the wide dynamic range of protein abundances in blood samples. Recent circulating blood proteome studies based on MS^11–13^ have achieved early successes also in disease biomarker discovery and understanding the pathogenesis of disease onset and progression^1–3^. The protein groups detected from blood using MS have been increased from about 300^1, 2, 14^ to over 1000 using high-abundance protein depletion kits^15^ and acid-assisted protein depletion^16, 17^.

In recent years, multiple nanoparticles (NPs) and MS-based methods further increased the protein identification to multiple thousand^18–21^. Selected NPs selectively adsorb circulating plasma proteins through non-covalent interactions (*e.g*., electrostatic, hydrophobic forces), forming a stratified protein corona categorized into hard and soft layers^22^. The “soft” corona, initially dominated by high-abundance proteins, is progressively replaced by a “hard” corona of high-affinity proteins through competitive displacement over time, as described by the Vroman effect^23^. The hard corona comprises tightly bound proteins with slow exchange rates, whereas the soft corona consists of loosely associated proteins in dynamic equilibrium with the surrounding environment, conferring distinct stability profiles under physiological conditions^24^. Beyond the application of diverse NPs in blood proteomics, emerging studies have shown that introducing small molecules to modulate NP protein corona diversity can achieve enhanced enrichment of low-abundance proteins^25–27^. Increasing number of NPs have been developed to measure circulating blood proteomes, reporting very different numbers of protein groups, ranging from ca 2000 to ca 7000, even using similar LC and MS instruments and methodologies. The variability of NP-based proteomic depth presents a major concern in this emerging field^28^.

A meta-analysis of 210 blood proteomics studies^29^ corroborates this premise, identifying contamination markers—including platelets, erythrocytes, or coagulation factors—in 54% (113/210) of analyzed studies. This study also has reported higher protein identification in platelets and erythrocytes compared to plasma samples, systematic investigations into how increasing platelet/erythrocyte contamination levels affect proteomic quantification and qualification remain lacking. More importantly, the impacts of such contamination in NP-enriched plasma proteomics workflows have not been thoroughly analyzed. We hypothesize that the presence of platelet and erythrocyte contaminants in plasma samples introduces variability in protein identification, with disproportionate impacts on low-abundance biomarkers.

To test whether platelet and erythrocyte contaminations are the major sources accounting for the observed huge variability of NP-based plasma proteomics experiments, we optimized a NP-based plasma proteome enrichment workflow called OmniProt capable of identifying ca 3000–6000 protein groups per 24-min gradient Astral nDIA^30^ analysis. We also generated a comprehensive spectral library of over 10,000 proteins detected in human plasma using more than 20 types of NPs. Building upon this analytical framework, we systematically evaluated the impact of platelet and erythrocyte contamination using the NP-based plasma proteomic workflow. With varying degrees of contamination, we identified signature protein biomarkers for each type of contamination. Furthermore, we developed an open-access software tool called Baize to evaluate NP-based plasma contamination. Finally, we evaluated the applications of OmniProt and Baize, in a cohort of 193 individuals, including 42 cases of pulmonary benign nodules and 114 small malignant nodules that conventional CT imaging cannot reliably distinguish the malignant tumor from the benign, necessitating surgical resection for definitive diagnosis.

## Results

### Selection and characterization of nanoparticles

To enrich low-abundance plasma proteins, we optimized our sample preparation using various silica-based NPs (**Fig. 1a**). Given that NP-mediated protein adsorption is largely non-specific and influenced by the physicochemical properties of both proteins and NPs^31^, we tested a panel of SiO\x{FFFF} NPs with distinct morphologies and sizes (**Supplementary Table 1**). This panel included solid spherical NPs (NP23 500 nm; NP24, 300 nm, NP25, 800 nm; and NP26, 1000 nm), mesoporous NP (NP84, 100–400 nm), hollow mesoporous NPs (NP85, ∼400 nm), and hierarchical porous NPs (NP86, 400–900 nm). To evaluate their performance in plasma proteome analysis, pooled plasma samples from lung cancer patients were analyzed in triplicate (15 µL per sample). Overall, over 80% of the total proteins identified by these NPs are shared by different NPs (**Supplementary Fig. 1a**). The reproducibility based on biological replicates is high, with coefficients of variation (CVs) of 1–2% (**Supplementary Fig. 1b**). These data suggest a high degree of technical reproducibility of this workflow. Solid silica NPs showed superior performance, yielding 16% more protein groups and 33% more peptides than other variants (**Fig. 1b-c**). Next, we further examined the solid silica microspheres of different sizes (300, 500, 800 and 1000 nm) using transmission electron microscopy (TEM) before and after binding plasma proteins (**Fig. 1d**). The TEM picture revealed a nearly uniform size for each type and monodispersed of solid silica NPs. After incubation with plasma samples, the diameters of all solid silica NPs increased, which is the so-called corona, in consistent with previously reported findings^32^. With the physical uniformity of the NPs established, we then evaluated their performance in enriching low-abundance proteins from plasma samples. Although the numbers of identified proteins and peptides were similar across the different sizes, practical considerations such as the specific surface area and the efficiency of washing by centrifugation to remove the protein corona favor medium-sized particles. Consequently, the 500 nm NPs (NP23) were selected as the optimal configuration for further characterization.

**Fig. 1.**
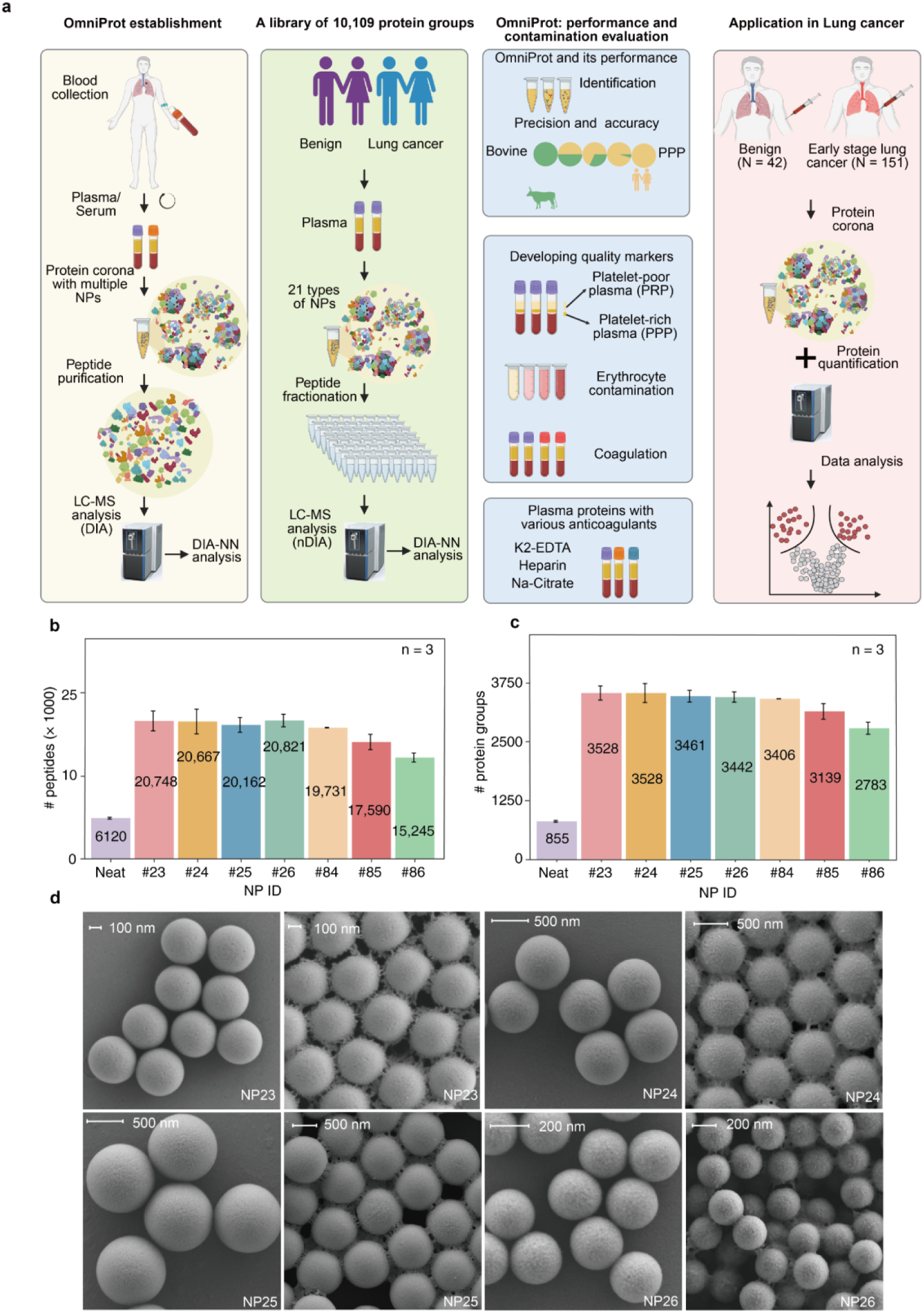
Systematic comparison of different SiO_2_-based nanoparticles. (a) Schematic overview of the study. Comparison of peptide (b) and protein group (c) identifications across SiO_2_-based NPs. (d) Morphological diversification of solid SiO\x{FFFF}-based nanoparticles revealed by transmission electron microscopy (TEM). Numerical labels denote cumulative identifications across triplicate analyses.

### Establishment of the OmniProt plasma proteomics workflow

NP-based protein enrichment^31, 33^ relies on protein corona formation and soft corona washing, both critical for capturing low-abundance plasma proteins. Using NP23, we optimized key procedures, including plasma dilution, protein corona formation, corona washing and NP storage. 15 μL aliquots of pooled plasma from lung cancer patients were analyzed using a 24-min gradient Astral nDIA analysis in triplicate. To optimize plasma dilution, we compared diluent buffers in acidic (3.0), neutral (7.0), and alkaline (11.0) conditions. Results showed that the alkaline diluent led to on average 36% more peptide identifications and 12% more protein identifications compared to the acidic and neutral conditions (**Supplementary Fig. 2a**). We further confirmed the preserved structural integrity of silica NPs and stable protein corona formation under the alkaline conditions using SEM analysis (**Fig. 1d**). Next, we optimized NP amount and incubation time. We tried different amounts of NPs ranging from 0.3 mg to 2.0 mg, and found that 0.5 mg slightly outperformed the other conditions (**Supplementary Fig. 2d**). We also incubated the NPs with the plasma for varying durations ranging from 30 min to 120 min, and found 30 min was sufficient (**Supplementary Fig. 2e**). Next, we evaluated the NP washing steps for removing soft corona, and found that three consecutive washes resulted in robust proteomics analysis (**Supplementary Fig. 2b**). We also compared different centrifugation speeds, and the results prioritized 7000 g as the optimal speed (**Supplementary Fig. 2c**). Finally, we assessed the thermal stability of NPs by incubating them at 4°C, 15°C, and 30°C for 24 hours. The results showed no significant difference in protein identifications, indicating that short-term exposure to room temperature does not affect NP performance (**Supplementary Fig. 2f**). These optimizations established the optimal conditions for protein corona formation, ensuring reliable and reproducible NP-based plasma proteome enrichment. We call this workflow as OmniProt in the following.

### Identification of over 10,000 protein groups from NP-enriched plasma proteomics

To construct a comprehensive spectral library for DIA data analysis in plasma proteomics, we utilized 21 chemically distinct NPs to enrich low-abundance proteins in pooled plasma samples from 20 patients with lung cancer. Notably, the plasma samples were absence of significant platelet, erythrocyte and coagulation-related contaminants. Peptide samples from these NP-enriched plasma proteins were fractionated into 30 or 60 fractions via high-pH reversed-phase chromatography (**Fig. 2a**), followed by nDIA Astral analysis. Altogether we generated 780 DIA files (**Supplementary Table 2**), which were then analyzed by DIA-NN, leading to identification of 126,661 peptide precursors mapping to 10,109 protein groups (**Fig. 2c**). The Human Plasma Atlas^34^ identified 4066 protein groups, of which 3819 protein groups (94%) are covered by our data resource (**Fig. 2c**). We then checked the proteins enriched by each NPs of various charge, functional group, matrix, hydrophobicity and the reaction class (**Fig. 2b**). The number of protein groups identified from the 21 NPs ranged from ∼2000 to ∼6000, with the lowest number observed for polystyrene NPs and the highest for silica-based NPs. Bioinformatics analysis of the 10,109 protein groups using Gene Ontology Biological Process^35, 36^ and KEGG^37^ databases showed predominant functional associations with signal transduction, protein phosphorylation, proteolytic regulation, apoptotic mechanisms, cellular adhesion and innate immunity (**Fig. 2d**). KEGG pathway analysis identified 207 significantly enriched pathways (FDR-adjusted p < 0.05), with the top decile comprising metabolic cascades, neurodegenerative disorders and oncogenic pathways (**Fig. 2e**), suggesting that this resource covers diverse physiological and pathological processes.

**Fig. 2.**
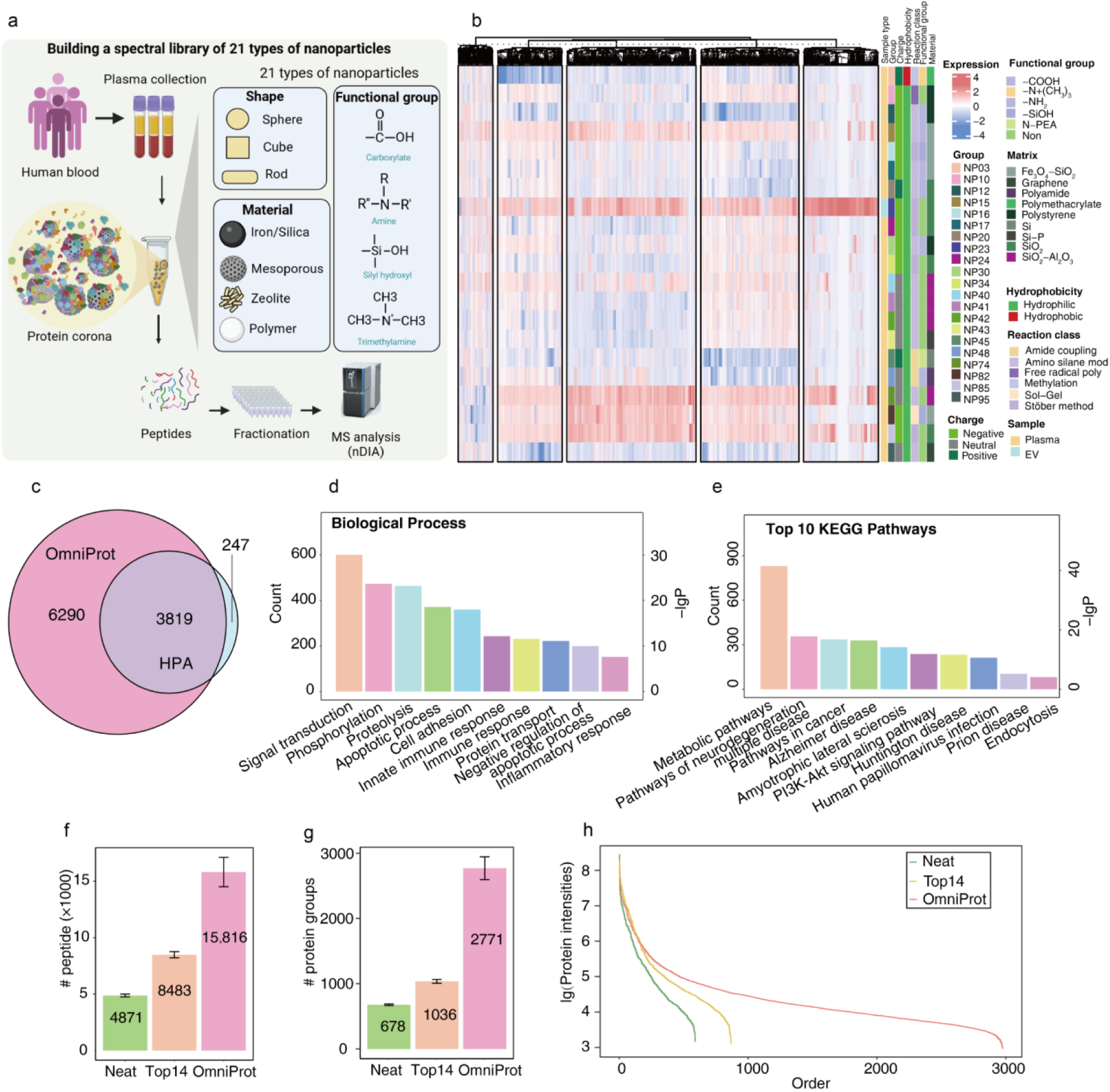
Nanoparticle-enhanced plasma spectral library construction and benchmarking. (a) Schematic view of nanoparticle-driven spectral library generation. (b) Heatmap overview of 10,109 protein groups identified across 21 nanoparticle types. (c) Comparative analysis with Human Plasma Atlas (HPA). Top 10 enriched KEGG pathways (d) and biological processes (e) ranked by DAVID GO analysis. Identified peptides (f), protein groups (g) and protein abundance ranking (h) in plasma samples processed with neat, Top14-depletion, and OmniProt methods.

### Performance of OmniProt in measuring low-abundance plasma proteins

Next, we employed this spectral library to perform a comparative analysis of OmniProt against both a commercial Top14 high-abundance protein depletion kit (Top14) and direct digestion of plasma samples (Neat). The evaluation utilized pooled platelet-poor plasma (PPP) samples analyzed by nDIA in an Astral. Our results showed that the OmniProt method identified a median of 2771 proteins and 15,816 peptides, yielding 2.6- and 5.7-fold more peptides (**Fig. 2f**) and 1.7- and 4.8-fold more protein groups (**Fig. 2g**) compared to the Top14 and the Neat condition, respectively. Notably, compared to Neat and Top14, OmniProt identified a greater number of lower-abundance proteins, with the highest concentration observed in the log_10_ intensity range of 3.2 to 4.5 (**Fig. 2h**). The above results initially show the improved sensitivity of the OmniProt workflow in detecting low-abundance plasma proteins. The lower identification is mainly caused by plasma quality issues from long-term frozen storage and repeated freeze-thaw cycles. In later experiments with newly frozen plasma samples, the OmniProt method successfully detected 2500-4200 protein group per plasma sample (**Fig. 7b**).

Next, to evaluate the precision of OmniProt-based proteome analysis, we designed a mixed-species benchmarking experiment^38^. We spiked increasing amounts of bovine plasma samples (0% to 100% volume-to-volume ratio) into a pool of human plasma samples. The mixed plasma samples were either analyzed by nDIA in an Astral or enriched for low-abundance proteins using OmniProt followed by nDIA analysis (**Fig. 3a**). The OmniProt samples led to identification of 36,977 peptide precursors derived from 4950 protein groups across all dilution conditions, with an 8-fold increase in protein identification compared to the neat sample (**Fig. 3b-c**). The OmniProt samples also showed comparable quantitative reproducibility to the neat samples across all spike-in ratios, with biological triplicates exhibiting median CVs < 20% (**Fig. 3d**). To assess accuracy, we focused exclusively on bovine-specific peptide precursors to eliminate interference from shared peptide precursors between the two species. This methodology systematically employs a broad bovine concentration gradient series to enable rigorous benchmarking of experimental fold-change measurements against theoretical under six predefined spiked-in ratio conditions. Results indicated that quantitative accuracy in multiple species samples remained generally consistent across dilution levels for both the neat and the OmniProt samples (**Fig. 3e**). Overall, these results show superior NP performance yielding more proteins than the neat plasma, and high degree of quantitative precision and accuracy of the NP-based plasma proteomics.

**Fig. 3.**
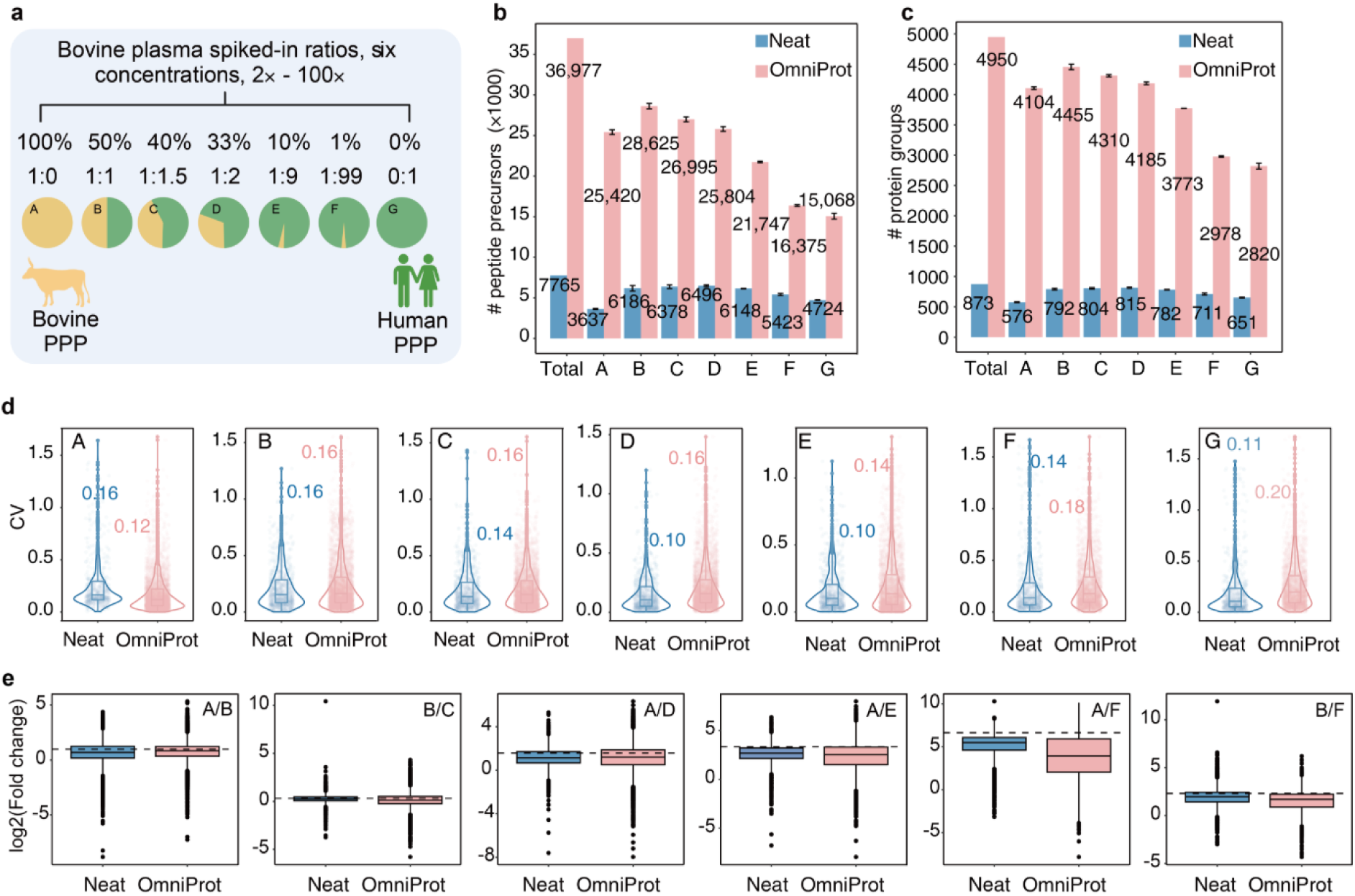
Benchmarking of OmniProt workflow. (a) Experimental design for spiking bovine plasma proteome into human plasma. Number of identified peptide precursors (b) and protein groups (c) in bovine-human plasma mixture. (d) Inter-species coefficients of variation (CVs) across spiked-in ratios. (e) Experimental fold change distributions versus theoretical expectations (dashed lines).

### Variability of protein identification in different plasma samples

During the optimization and applications of the OmniProt workflow, we observed that this protocol, when applied to different plasma samples, led to a largely variable number of protein identifications, ranging from ca 3000 to 7000 (**Supplementary** Figure 3a-d). Then we inspected the data, and found that the samples with high number of protein identifications contained abnormally high levels of platelet-derived and/or erythrocyte-derived proteins, suggesting potential contamination of platelets and erythrocytes, which was subsequently thoroughly investigated as described below.

### Platelet contamination and markers

To systematically assess platelet contamination in NP workflows, we collected platelet-rich plasma (PRP) and the PPP samples from two donors (one male and one female) and pooled the respective PRP and PPP samples from each donor individually. Subsequently, the PRP and the PPP samples were mixed at seven pre-defined volumetric ratios (**Fig. 4a**). The mixed samples were processed with the neat plasma and the OmniProt workflow in triplicate, followed by nDIA analysis (**Fig. 4a**). Our results showed that in both the neat and the OmniProt samples, platelet spike-in led to increased identification of peptide precursors and protein groups (**Fig. 4b-c**). The OmniProt workflow identified on average 4580 protein groups in the PRP samples, while only 2492 protein groups in the PPP samples.

**Figure 4.**
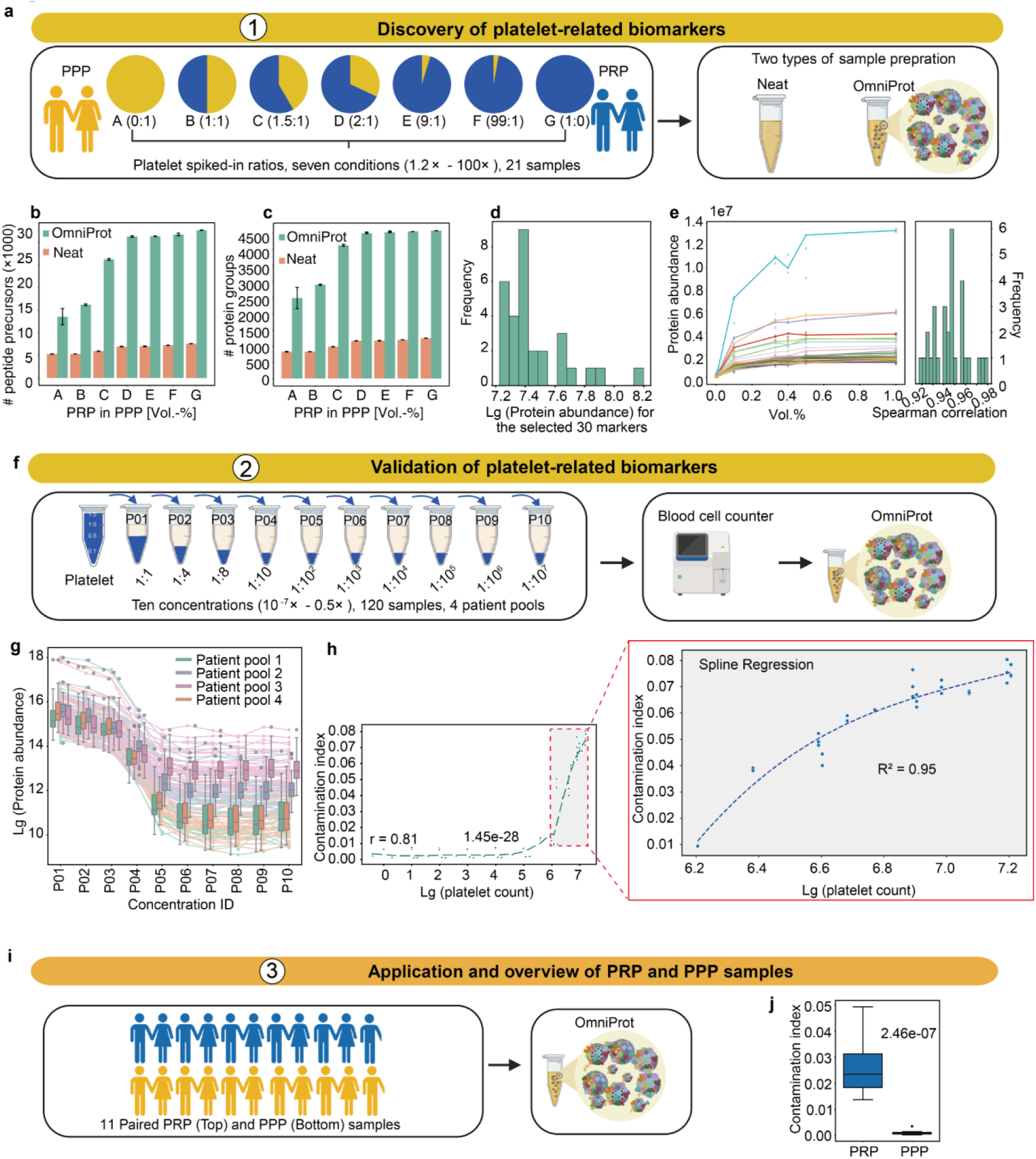
Quality marker panel for platelet contamination. (a) Experimental workflow for platelet-related biomarker discovery. Number of identified peptide precursors (b) and protein groups (c) in discovery dataset. (d) Abundance distribution profiles of 30 selected protein biomarkers. (e) Spearman correlation analysis of platelet-related biomarkers in discovery dataset. (f) Validation workflow of platelet-related biomarkers. (g) Z-scored protein intensities of 30 platelet-related biomarkers across dilution series. (h) Correlation between platelet count and contamination index. (i) Application scheme for platelet contamination assessment. (j) Contamination index calculation in the PRP and PPP samples. PRP: platelet-rich plasma, PPP: platelet-poor plasma.

Next, we asked whether platelet contamination alters the concentration of circulating plasma proteins after NP enrichment. We identified 2432 proteins in pure PPP samples with no significant association with platelet levels (correlation coefficient < 0.7). These proteins are regarded as platelet-independent circulating proteins to a certain extent. Our data showed that the abundances of these proteins in the PPP samples exhibited relatively weak correlation (r = 0.54–0.73) with those in plasma samples with various degrees of platelet contamination (**Supplementary** Figure 4a). This indicates that NP-based enrichment moderately alters these platelet-independent circulating proteins, thereby compromising the quantitative accuracy of circulating plasma proteome analysis. Thus, it is non-trivial to assess the platelet contamination in NP-based plasma proteome analysis.

Geyer^29^ *et al* have identified 30 proteins as biomarkers for platelet contamination in neat plasma proteome analysis. Next, we evaluated whether these protein markers could be applied to evaluate platelet contamination in the above-mentioned neat and OmniProt samples. As expected, the neat samples showed strong correlation (median r = 0.95) across 30 protein markers, confirming the findings from Geyer^29^ *et al*. However, the OmniProt samples showed relatively weaker correlation (median r = 0.75) as shown in **Supplementary** Figure 4b-c. The protein GSN even showed negative correlation with increasing platelet contamination. Therefore, there is a need to further define a customized contamination list for NP-based plasma proteome experiments.

To identify protein biomarkers for monitoring platelet contamination, we first filtered out all the protein groups with missing values exceeding 50% across all samples, leading to 4404 consistently detected protein groups, which were subsequently clustered into eight distinct groups using Mfuzz. Mfuzz clustering divided the proteins into two groups with different abundance trends. The proteins in the first group gradually increased with platelet concentrations until plateau, while the proteins in the other group declined as platelet levels increased (**Supplementary** Figure 5a). We then identified the top 100 proteins with highest correlation with increased platelet concentrations. From this subset, the top 30 proteins exhibiting highest abundance levels were retained as candidate biomarkers for platelet contamination. The protein expression of these 30 biomarkers in plasma samples with increasing platelet contamination is provided, with Spearman correlation coefficients > 0.92 (**Supplementary** Figure 5c), confirming their correlation with platelet contamination. These proteins are of relatively high abundance in the NP-enriched samples (**Fig. 4d**). The Spearman correlation of our markers showed stronger consistency (median = 0.95, range = 0.92–0.98) (**Fig. 4e**). The selected platelet-associated proteins, including ADP/ATP translocase 2 (SLC25A5)^39^, platelet factor 4 variant 1 (PF4V1)^40^, thromboxane A synthase 1 (TBXAS1) and integrin beta-1 (ITGB1), exhibit critical roles in platelet activation and aggregation pathways, substantiating their value as contamination biomarkers. Pathway enrichment analysis revealed that these 30 proteins are primarily associated with mitochondrial protein degradation, mitochondrial calcium ion transport, eicosanoid signaling, sirtuin signaling pathway and cardiac hypertrophy signaling (Enhanced) (**Supplementary** Figure 5b).

To further validate these 30 biomarkers, pure platelet and PPP samples from 12 patients were procured and pooled into four groups (three patients per group). These platelet samples were subjected to a ten-step series dilution with PPP samples. We further performed blood cell counting to confirm the degree of platelet contamination. These samples were analyzed with the OmniProt workflow (**Fig. 4f, Supplementary Table 3**). The results confirmed that higher degree of platelet contamination led to increased number of peptide precursor and protein group identification (**Supplementary Fig. 6**. We also confirmed that the abundance levels of the 30 platelet markers reliably reflect the degree of platelet contamination (**Fig. 4g**).

Next, we developed a computer program called Baize (https://www.guomics.com/Baize) to evaluate platelet contamination by calculating the ratio of summed signals from 30 platelet markers to total protein intensity, followed by spline regression to predict platelet levels in plasma samples. The model achieved an R² of 0.95, showing strong correlation for systematic platelet contamination assessment in NP-based plasma proteomes (**Fig. 4h**). The platelet contamination algorithm was further applied to paired PRP and PPP samples obtained from 11 patients with lung cancer (**Fig. 4i**). The results showed that the degree of platelet contamination in the PPP samples remained close to zero, while that in the PRP samples ranged from 0.15 to 0.48 (median: 0.22, **Fig. 4j**). Indeed, the abundance of these 30 proteins is significantly different between the PPP and PRP samples (**Supplementary Fig. 7**. The platelet contamination also influenced the proteome profiling. The neat samples led to identification of 955 protein groups in the PPP samples and 1373 in the PRP samples, while the OmniProt method improved the protein group identification to 2701 in the PPP samples and 4328 in the PRP samples. 94.5% of the proteins identified in the neat samples were also identified by the OmniProt method (**Supplementary Fig. 8a-b**). We performed hierarchical clustering analysis on proteins identified in both OmniProt- and neat-processed PPP and PRP samples. The analysis revealed that compared to neat samples, the proteins uniquely identified in OmniProt samples are primarily distributed in the following lung cell types: Travaglini lung trem2 dendritic cell, Descartes fetal lung myeloid cells, Travaglini lung platelet megakaryocyte cell, and Travaglini lung proximal basal cell. In the OmniProt-processed samples, the PRP samples exhibited elevated levels of proteins linked to lung cell lineages compared to the PPP samples, specifically Travaglini lung platelet megakaryocyte cell and Travaglini lung trem2 dendritic cell (**Supplementary Fig. 8c**).

Finally, we asked whether these biomarkers could be used for the samples prepared by other types of nanoparticles. We selected two frequently reported nanoparticle types: Zeolite NaY^33^ (NP74) and silanol-functionalized Fe_3_ O_4_ ^19^(NP81). We collected the blood sample from a donor and prepared PPP and PRP plasma samples. 100 µL of PPP and PRP plasma samples were pooled and subjected to a ten-step serial dilution. The diluted plasma samples were processed using the two nanoparticles, namely NP74 and NP81, followed by nDIA analysis. Our results confirmed that increasing platelet concentration elevated peptide precursor and protein group identifications for both NP74- and NP81-processed plasma samples, reaching over 6000 protein groups per injection (**Supplementary Fig. 9a-d**). The 30 platelet-related biomarkers showed a high degree of correlation with increasing platelet contamination in NP74-(median: 0.95, range 0.89–0.96) and NP81-processed samples (median: 0.93, range 0.87–0.94). Thus, our results further support the applicability of the platelet-related biomarkers identified in this study for evaluating platelet contamination in different NPs (**Supplementary Fig. 9e-f**). Notably, the samples with identical platelet contamination levels showed different degrees of contamination between NP74- and NP81-processed samples, ranging from 0.000 to 0.055 for NP81 and 0.000 to 0.035 for NP74 (**Supplementary Fig. 9g–h**). These findings indicated that the degree of platelet contamination cannot be directly compared between samples processed through different NP workflows.

### Erythrocyte contamination and markers

Hemolysis represents another critical preanalytical variable in plasma proteomics. To systematically assess erythrocyte contamination in NP-based plasma experiments, we applied similar strategy to the platelet contamination algorithm for systematic analysis. We collected pure erythrocytes and the PPP plasma samples from two donors (one male and one female) and pooled the respective erythrocyte and PPP plasma samples from each donor separately. Then, we mixed the erythrocytes and PPP plasma samples with six pre-defined volumetric ratios (**Fig. 5a**). The mixed samples were processed with the neat plasma and the OmniProt workflow in triplicate, followed by nDIA analysis (**Fig. 5a**). Our results showed that in both neat and OmniProt samples, erythrocyte spike-in led to increased identification of peptide precursors and protein groups, except for the 100% RBC samples (**Fig. 5b-c**). Next, we investigated whether erythrocyte contamination affects circulating plasma protein concentrations following OmniProt enrichment. We identified 986 proteins in the PPP samples showing no significant association with erythrocyte levels (Spearman correlation coefficient r < 0.7). Likewise, these proteins are regarded as erythrocyte-independent circulating proteins to certain extent. Our data revealed weak correlations (r = 0.59–0.93) between the abundance of these proteins in the PPP and the erythrocyte-contaminated plasma samples (**Supplementary Fig. 10a**). These observations show that NP-based enrichment perturbs even erythrocyte-independent circulating proteins, ultimately compromising the quantitative accuracy of circulating plasma proteome analysis. To further evaluate the abundance distribution of erythrocyte-independent circulating proteins and circulating plasma proteins, we categorized protein distributions in samples with various erythrocyte contamination levels through Mfuzz clustering analysis, revealing four clusters of proteins elevated with increasing RBC contamination, while another four clusters showed reverse pattern (**Supplementary Fig. 10b**).

**Figure 5.**
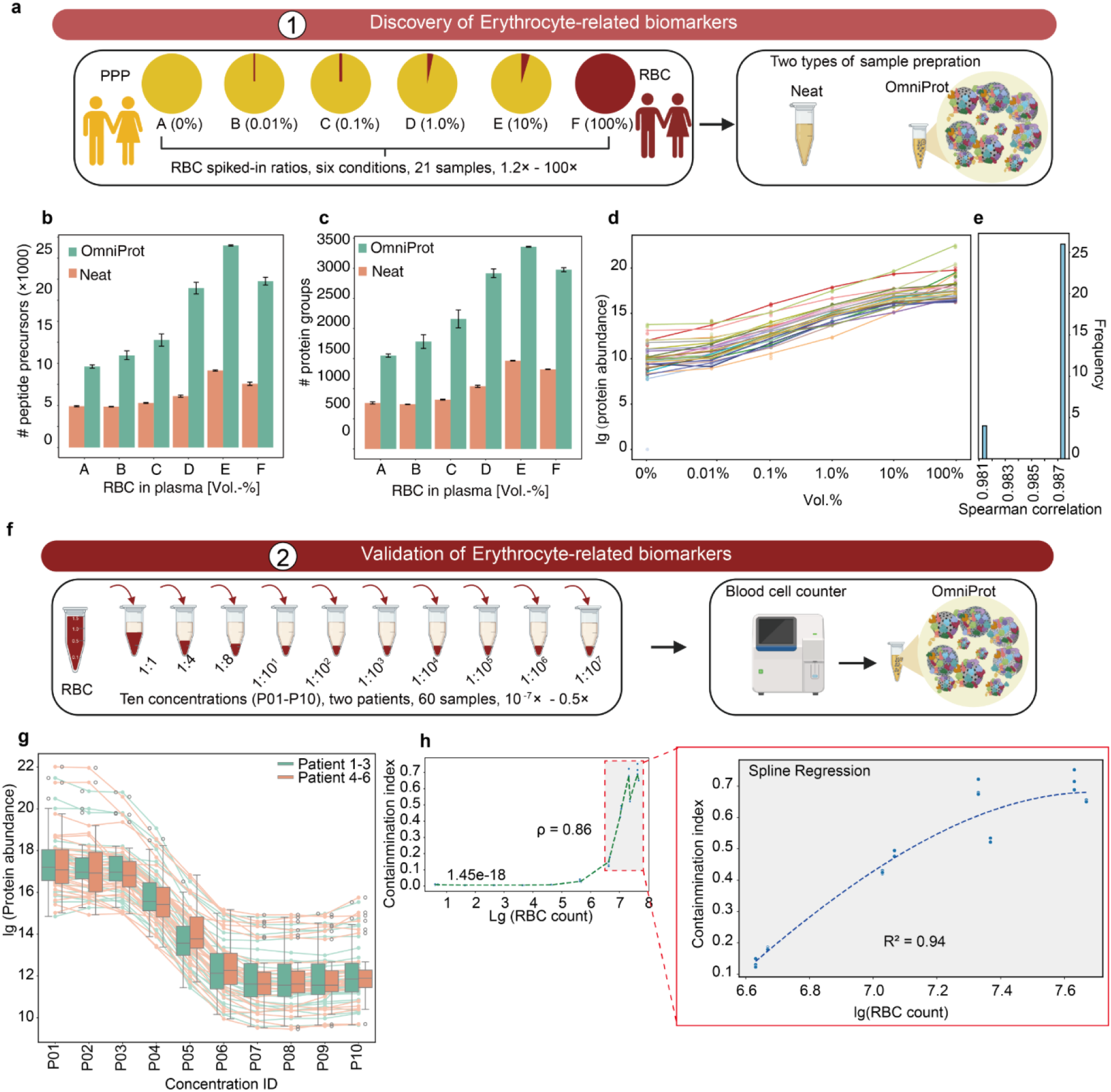
Quality marker panel for erythrocyte contamination. (a) Discovery workflow for erythrocyte-related markers. Number of identified peptide precursors (b) and protein groups (c) in discovery dataset. (d) Spearman correlation analysis of 30 erythrocyte-related biomarkers in discovery dataset. (e) Abundance distribution profiles of erythrocyte-associated markers. (f) Validation workflow for erythrocyte contamination markers. (g) Z-scored protein intensities of erythrocyte markers across dilution series vs. spike-in proportion. (h) Correlation of erythrocyte count and contamination index. PPP: platelet-poor plasma.

Geyer^29^ *et al* have identified 30 proteins as biomarkers for erythrocyte contamination in neat plasma proteome analysis. Here, we evaluated whether these protein markers could be applied to evaluate erythrocyte contamination in the above-mentioned neat and OmniProt samples. As expected, the neat samples showed strong correlation (r = 0.98–1.00) across 30 protein markers, confirming the findings from Geyer^29^ *et al*. However, the OmniProt samples showed relatively weaker correlation (r = 0.75–0.98) as shown in **Supplementary** Figure 4d-e. To identify protein biomarkers for monitoring RBC contamination, we first selected 100 proteins showing the strongest correlation with RBC levels, then selected the top 30 candidates based on protein intensity for RBC contamination assessment. The 30 candidate proteins exhibited robust correlation (**Fig. 5d-e, Supplementary Fig. 11a-b**). These proteins were rarely detected in the plasma samples without RBC contamination but showed higher intensities in the 10% RBC samples (**Supplementary Fig. 12a**). Pathway enrichment analysis revealed that these markers are primarily associated with erythrocyte gas exchange (oxygen uptake/carbon dioxide release and *vice versa*), neutrophil degranulation, JAK-STAT signaling-mediated gene/protein expression following IL-12 stimulation, and COPI-mediated anterograde transport (**Supplementary Fig. 11c**). We then evaluated the analytical performance of our 30 newly identified biomarkers in those samples. The Spearman correlation of our markers showed stronger consistency (median r = 0.987) (**Fig. 5e**), including RBC-specific markers with distinct biological roles. Hemoglobin subunits^34^ (HBD, HBB, HBA1), essential for oxygen transport, showed strong RBC specificity. The membrane-stabilizing spectrins^41^ (SPTA1, SPTB) maintain RBC structural integrity, while SLC4A1^42^ regulates gas exchange through anion transport.

To further validate these biomarkers, the erythrocyte and the PPP samples from six patients were collected and pooled into two groups (three patients per group). These erythrocyte samples were subjected to a ten-step serial dilution with PPP samples (**Fig. 5f**). We further performed blood cell counting to confirm the degree of erythrocyte contamination. These samples were analyzed with the OmniProt workflow (**Supplementary Table 3, Fig. 5f**). The results confirmed that higher degree of erythrocyte contamination led to increased number of peptide precursor and protein group identification (**Supplementary Fig. 12b**). We also confirmed that the abundance levels of the 30 erythrocyte markers reliably reflect the degree of erythrocyte contamination (**Fig. 5g**). Next, we developed erythrocyte contamination indices following established methodologies previously applied to platelet contamination assessment and implemented in the Baize software. This novel metric exhibited remarkable robustness, showing strong linear correlation (R² = 0.94) with cell counts obtained by hematocytometer (**Fig. 5h**).

### Variability introduced by coagulation

Improper blood collection represents a critical yet often overlooked source of variability in plasma proteomics. Proper collection of plasma samples requires immediate gentle inversion after blood withdrawal to ensure uniform mixing of anticoagulants, followed by centrifugation to yield cell-free plasma. Delays in this process risk partial coagulation, creating hybrid plasma-serum matrices that distort biomarker quantification. Plasma samples are typically collected using anticoagulant-containing tubes: EDTA (purple-top), sodium citrate (blue-top), or lithium/sodium heparin (green-top). All the tubes are used in various clinical applications; however, no systematic study has investigated their impacts in NP-based plasma proteomics.

To evaluate the impact of anticoagulants on plasma proteomics, we procured plasma samples from 10 donors (5 males, 5 females). Each donor contributed triplicate blood samples using the three kinds of anticoagulation tubes. Both neat plasma and OmniProt-processed plasma samples were analyzed by nDIA MS. Our results showed that the plasma samples processed via neat and OmniProt showed no marked differences in the total numbers of peptide and protein identification across the three tube types (**Fig. 6a**). The neat samples led to the identification of 1371-1468 protein groups, while the OmniProt samples yielded 4668 proteins (**Fig. 6a**). However, only 59.2% of the proteins identified in the OmniProt samples are shared across all tubes (**Fig. 6b**), suggesting the proteome variability of plasma samples collected in different tubes. Compared to the neat samples, the OmniProt samples exhibited reduced protein signals associated with estrogen receptor signaling, mitochondrial dysfunction and related protein degradation and sirtuin pathways. Conversely, enhanced enrichment was observed for proteins involved in ROBO receptor signaling, ribosomal quality control, coronavirus replication machinery, eukaryotic translation initiation, B cell developmental processes, FcRIIB and PI3K signaling in B lymphocytes, as well as IL-15 and p70S6K pathways (**Fig. 6c**). OmniProt-processed EDTA and heparin plasma samples exhibited greater similarity in protein enrichment profiles compared to other tube types, with better enrichment of proteins associated with B cell development, FcRIIB and PI3K signaling in B lymphocytes, IL-15 signaling and p70S6K pathway activation.

**Figure 6.**
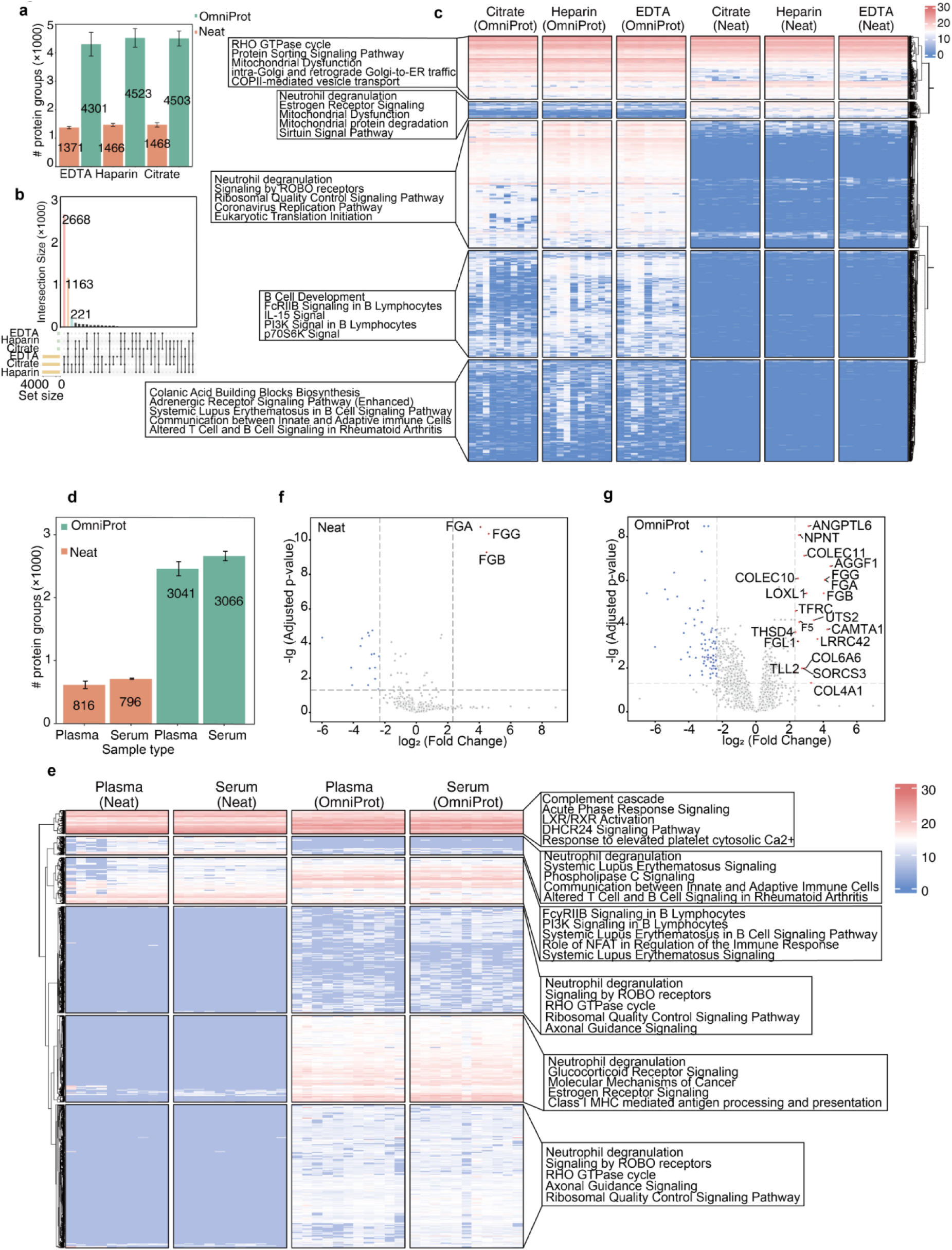
Performance evaluation of OmniProt in different blood collection tubes. (a) Protein group identification yields in three blood collection tube types. (b) Upset plot of protein overlap across three blood collection tube types. (c) Heatmap visualization of tube-specific protein profiles. (d) Protein identifications from 10 paired plasma and serum samples. (e) Heatmap overview of paired plasma/serum proteomes. Volcano plots of plasma vs. serum samples processed with neat (f) and OmniProt (g).

We also evaluated the impact of partial coagulation which results in hybrid plasma-serum matrices. Paired plasma and serum samples were collected from 10 donors (5 males and 5 females) using EDTA-coated and silica-coated blood collection tubes, respectively. Each donor provided two blood samples that were analyzed as neat samples and the OmniProt workflow. OmniProt proteomic profiling identified 3041 protein groups in the plasma and 3066 in the serum samples, surpassing the neat plasma and serum samples (816 and 796 proteins respectively) (**Fig. 6d**). Heatmap analysis showed that OmniProt-exclusive proteins predominantly mapped to neutrophil degranulation pathways, systemic lupus erythematosus signaling cascades, and ROBO receptor-mediated signaling networks (**Fig. 6e**). To prioritize coagulation-related proteins, we selected the proteins through following criteria, requiring a minimum fold-change of 5 and statistical significance (adjusted p-value < 0.05) in both processing methods (**Fig. 6f-g**). These differentially expressed proteins contain FGA, FGG, and FGB which are markedly upregulated in the neat samples, consistent with prior literature^29^. The OmniProt samples additionally highlighted 17 known coagulation-related proteins including AGGF1 and TFRC. Finally, we developed a method to evaluate coagulation-related contamination based on these 20 proteins and implemented it in Baize.

### Baize software for evaluating contamination in NP-based plasma proteomics

Here, we introduce Baize, a web-based software tool for rapid evaluation of sample contamination in three critical dimensions: platelet contamination, erythrocyte lysis, and residual coagulation protein carryover. The algorithm calculates cell-type-specific contamination index by normalizing the summed intensity of marker proteins against the total intensity of plasma proteome (Contamination Index = Marker Protein Intensities / Σ All Plasma Protein Intensities). Users can submit a protein matrix, and then Baize will output an evaluation of contamination for every plasma sample. Baize is freely accessible at https://www.guomics.com/Baize.

### Application in a lung cancer cohort

To evaluate the application of OmniProt and Baize, we procured 193 individuals including 42 with benign nodules, and 151 with early-stage malignancies. In these cases, conventional CT imaging often produces inconclusive results, necessitating surgical resection for definitive diagnosis. We also included six patients with advanced-stage lung cancers as positive controls. Detailed clinical characteristics of the cohort are summarized in **Supplementary Table 4** and **Fig. 7a**. Plasma specimens were collected as PPP and processed in seven randomized batches using the OmniProt workflow followed by nDIA analysis. We also included seven randomly selected biological replicates and seven randomly selected technical replicates to assess reproducibility. The proteomics analysis led to identification of a median of 4413 protein groups per plasma sample (**Fig. 7b**) with stable CVs observed across both biological and technical replicates (**Fig. 7c**). The potential contamination of these samples for platelets, erythrocytes, and coagulation factors were assessed via Baize software. Results showed five samples exhibited detectable platelet, erythrocyte or coagulation contamination (**Fig. 7d-f**). Samples with contamination were excluded from downstream analysis.

**Figure 7.**
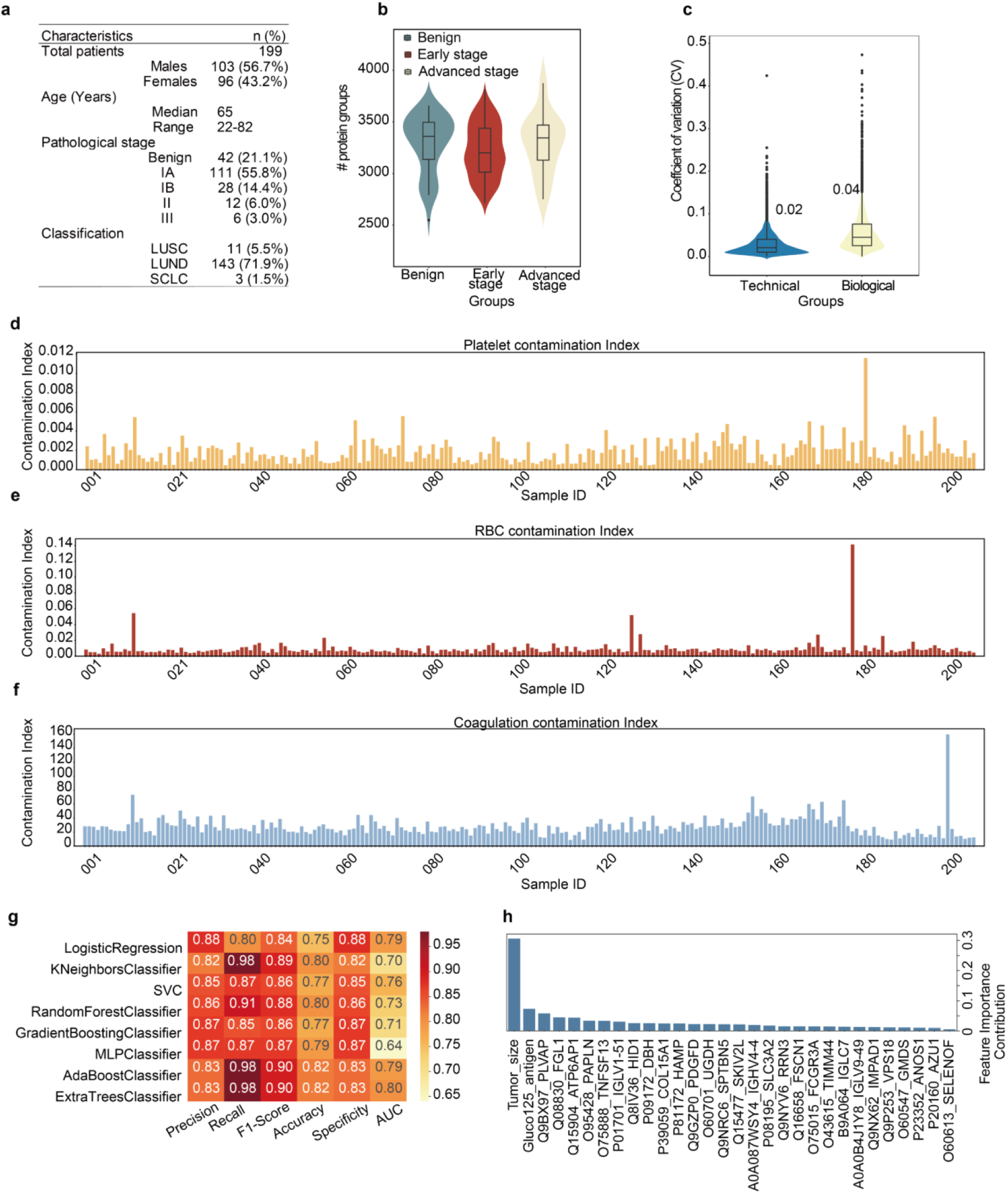
OmniProt-enabled plasma proteomics facilitates early detection of lung cancer. (a) Clinical characteristics of lung cancer cohort. (b) Global distribution of identified protein groups in the plasma proteome. (c) Reproducibility analysis in technical and biological replicates. Platelet-(d), erythrocyte-(e) and coagulation-(f) contamination profiling in the lung cancer cohort. (g) Machine learning classifier performance comparison. (h) Feature importance ranking of diagnostic biomarkers.

We then applied eight machine learning algorithms, including logistic regression, kNeighbors, SVC, Random Forest, GradientBoost, MLP, AdaBoost and Extra Trees, to separate benign and malignant lung nodules (**Fig. 7g**). Top 30 proteins or clinical features were selected for modeling. The results showed that the Extra Tree has the highest F1 score, accuracy, precision, recall, and AUC values compared to the other models (**Fig. 7g**). The Extra Tree model prioritized tumor size, CA-125 and 28 proteins with known association with lung cancer, including Fascin-1 (FSCN1)^43^, CD98hc (SCL3A2)^44^, Platelet-derived growth factor D (PDGF)^45^, CD16a (FCGR3A) ^46^, Hepcidin (HAMP)^47, 48^, Azurocidin 1 (AZU1) and Collagen XV (COL15A1)^49^, suggesting that our methodology successfully identified critical proteins related to the disease phenotype. Further validation of these potential protein biomarkers in independent lung nodule cohorts is beyond the scope of this study.

## Discussion

Circulating blood proteomics has become an indispensable strategy for biomarker discovery, due to its minimally invasive sampling approach and the capacity to systematically characterize disease-associated molecular signatures within blood components^1–3, 12, 50^. MS-based plasma proteomics analysis faces challenges in proteome coverage due to the wide dynamic range of plasma protein abundances. Various technological advancements — such as high-abundance protein depletion kits^15^, acid precipitation^17, 51^, and recently emerged nanoparticle enrichment methods^19–21, 52^ — aim to improve the depth of plasma protein identification. The nanoparticle-based approach is gaining popularity, but both previous literatures^19, 20, 33^ and our current study report significant variability in protein identification results (ca 2000–7000 protein groups) across plasma samples even with similar LC-MS workflows. Based on prior evidence, we hypothesized that platelet and erythrocyte contamination might contribute to this variability. In this study, we established OmniProt – a deep-coverage nanoparticle-enhanced plasma proteome preparation workflow – accompanied by the Baize computational toolkit, which enables rapid assessment of platelet, erythrocyte, and coagulation-related contamination in clinical plasma samples. This advancement carries implications for several key contributions for the plasma proteome community.

The first contribution of this study is that we confirm that the variability of circulating plasma protein numbers identified by nanobeads-based methods is mainly due to contaminations of platelet, erythrocyte and coagulation. In our spike-in experiments, the protein numbers identified from plasma increase sharply when the contamination of platelet and erythrocyte deteriorates. Although such observations have been reported for neat plasma by Geyer *et al*^29^, no study has comprehensively investigated the impacts of contamination in nanoparticle-based circulating plasma proteomics.

After realizing the impact of blood contamination in plasma protein identification, our next contribution is to clarify that such contaminations influence protein quantification. Our data showed that the abundances of circulating plasma proteins in the PPP samples exhibited weak correlation (r = 0.54-0.73) with those in plasma samples with various degrees of platelet contamination.

We further developed a software tool called Baize to evaluate contamination in nanoparticle-based circulating plasma proteomics experiments. We discovered and validated protein biomarkers for evaluating contaminations of platelet, erythrocyte, and coagulation-related proteins. A spline regression algorithm was subsequently implemented to assess contamination severity and quantify contamination levels. The evaluation is implemented in an open-access web-based software tool named Baize (https://www.guomics.com/Baize).

In this study, we also established a silica nanoparticle-based sample preparation workflow for enriching low-abundance plasma proteins. From previous studies, multiple nanoparticle enrichment methods have been used for plasma proteomics^19–21, 52^. Representative approaches include magnetic material-mediated low-abundance protein enrichment (5-10 types) as commercialized by Seer^19, 53, 54^, molecular sieve-based methods reported by Ma *et al*^20^ and Li *et al*^33^, and silica-based strategies (ProteoFish) developed by Tan *et al*^21, 52^. During our preliminary experiments, molecular sieve-type nanoparticles exhibited performance degradation in low-abundance protein enrichment following moisture absorption, with unrecoverable efficiency even after secondary drying – indicating potential risks for cross-laboratory and longitudinal large-cohort applications^55^. Silica-based NPs show high degree of stability, batch-to-batch consistency, and cost-effectiveness. Tan’s team achieved an average of ∼1174 protein group identifications using silica nanoparticle enrichment on an Exploris 480 instrument with a 120-min LC gradient method^21^. In this study, we systematically compared the effects of different types (morphologies, sizes) of silica nanoparticles on plasma proteome identification and selected NP23 (solid, 500 nm) with the highest identification capacity and stability for subsequent analyses. We then optimized key parameters in protein corona formation, including plasma dilution, incubation time, washing steps, and thermal stability, establishing an optimized workflow named OmniProt. OmniProt enables the identification of ca 3000–6000 proteins across different plasma samples using Astral with 60 SPD throughput, yielding 1.7- and 4.8-fold more protein groups and 2.6- and 5.7-fold more peptides compared to the Top14-protein depleted samples and the neat plasma samples, respectively. In the mixed-species benchmarking experiment, the OmniProt method exhibited high precision and high accuracy comparable to neat samples. Nevertheless, the silica beads require multiple centrifugation steps which introduce difficulties for automated sample preparation. Increasing magnetic bead-based nanoparticles have been developed for plasma proteomics^19, 53^. Magnetic version of OmniProt should be developed in future studies.

Our study also presents a comprehensive Astral spectral library for plasma proteomics. Several spectral libraries for plasma proteome have been published. Tan *et al*^21^. established a SiO_2_ nanoparticle-derived library covering 2564 proteins, whereas Ma *et al*. developed a Zeolite NaY nanoparticle-based database containing 6524 proteins^20^. In this study, we constructed a comprehensive spectral library for DIA data analysis using 21 chemically distinct nanoparticles combined with peptide fractionation, containing 126,661 peptide precursors corresponding to 10,109 protein groups. This spectral library was established using cell-free plasma samples devoid of cellular contamination and represents the first publicly available, extensively characterized plasma proteome spectral library for community use, as supported by our literature review.

Finally, we validate the OmniProt and Baize tools with a patient cohort of 193 individuals with CT-indistinct benign small pulmonary nodules or early-stage lung cancers. We identified on average ∼4000 proteins per plasma sample with OmniProt using Astral at 60 SPD throughput, with five samples flagged for platelet/erythrocyte or coagulation contamination. Furthermore, by integrating proteomic data with clinical information using machine learning, we developed a blood-based classifier to distinguish benign pulmonary nodules from early-stage lung cancers, achieving an AUC of 0.8. This result demonstrates that the OmniProt provides a robust pipeline for the development of multiplexed protein biomarker detection in lung cancer screening.

While Baize effectively identifies plasma samples with potential platelet, erythrocyte or coagulation-related contamination, our study has several limitations. First, Baize was developed using 6–10 groups of plasma samples with varying contamination levels; due to limited data volume, the current implementation can only assess contamination severity but cannot systematically correct for contamination-induced quantitative biases in proteomic analyses. Although OmniProt demonstrated high accuracy in distinguishing small malignant nodules from benign ones in our clinical cohort, a second limitation lies in the retrospective nature of the early lung cancer screening cohort comprising 193 patients. Future validation through prospective large-scale cohorts will be essential to refine biomarker panels and establish precise diagnostic or prognostic applications. However, this is beyond the scope of this current study.

In conclusion, we systematically investigated contamination in nanoparticle-based circulating plasma proteomics, and found that contamination of platelet, erythrocyte, and coagulation-related proteins significantly alters the number of protein identification and the protein quantities. We then identified protein biomarkers to evaluate these contaminations and present an open-access software Baize for evaluating the contamination of NP-based plasma proteomics.

## Materials and methods

### Characterization of Diverse SiO_2_-Based NPs

All NPs were purchased from Westlake Omics. The morphology of SiO_2_ NPs was analyzed using a Zeiss Gemini 550 Crossbeam (Oberkochen, Germany) and CIQTEK SEM5000X FESEM Microscope (Anhui, China) with gold-coated copper TEM grids. For SEM sample preparation, both NPs and NP-protein coronas were first fixed overnight at 4°C in a fixative solution (pH 7.2) containing 2% paraformaldehyde (PFA) and 2.5% glutaraldehyde. After fixation, the samples were washed three times with 0.1 M PBS at 4°C on a shaker for 15 min. Subsequently, the samples were rinsed three times with deionized water at room temperature on a shaker, followed by centrifugation at 7000 g for 3 min. After the final centrifugation, 100 µL of the supernatant was retained. The pellet and 100 µL supernatant were then sonicated for 3 min in an ice bath to yield a suspension. For SEM imaging, 2.5 µL of the suspension was dropped onto a silicon wafer and dried for 2 min using a drying system (Leica, Germany). The dried samples were then coated with an 8 nm layer of platinum using an EM ACE600 High Vacuum Sputter Coater (Leica, Germany). The platinum-coated samples were subsequently used for further analysis.

### Clinical sample collection

Blood samples were collected from patients with lung cancer or pulmonary benign nodules using standardized vacutainer tubes at the First Affiliated Hospital of Zhejiang University School of Medicine (Hangzhou, China), with ethical approval granted by both the Institutional Ethics Committee of the First Affiliated Hospital (Approval No. 2024-0511) and the Ethics Committee of Westlake University (Approval No. 20231218GTN001). Four types of blood collection tubes, including EDTA K2 (purple-top) and serum (red-top) tubes from Hengshui Shengcixing Company (Hebei, China), as well as lithium heparin (green-top) and lithium citrate (light blue-top) tubes from Gongdong Medical (Taizhou, China), were used to collect plasma and serum samples.

In the platelet contamination assessment experiment, we obtained four types of plasma samples from whole blood samples using four isolation protocols. The first protocol involved dual centrifugation cycles at 2000 g for 15 min each, collecting the plasma supernatant after the second centrifugation. The second protocol applied a single centrifugation step at 2000 g for 15 min, collecting the plasma supernatant. The third protocol utilized a lower centrifugal force of 200 g for 15 min, collecting the plasma supernatant. The fourth protocol employed overnight sedimentation followed by supernatant collection, collecting the plasma supernatant. All processed plasma samples were immediately aliquoted into 1.5 mL tubes and stored at −80°C. Prior to experimental use, frozen aliquots were thawed at 4°C under controlled conditions.

Whole blood samples collected for quality marker panel establishment were first analyzed for cellular components within 4 hours post-collection using an automated hematology analyzer (Shenzhen Mandray Biomedical Electronics Co., Ltd., China). Purified PRP, PPP, erythrocyte, and platelet samples were obtained through the following procedures. The blood was first centrifuged at 200 g for 10 min, retaining both the pellet and supernatant. To minimize erythrocyte contamination in the supernatant, approximately 0.5 cm of supernatant adjacent to the erythrocyte layer was discarded, resulting in PRP samples. The pellet was centrifuged at 2000 g for 15 min, and the upper layer (containing residual plasma, buffy coat, and ∼1 mL erythrocytes) was removed, yielding erythrocyte-enriched fractions. Subsequently, erythrocytes were washed twice with 6 mL of phosphate-buffered saline (PBS) containing 1.6 mg/mL EDTA, followed by centrifugation at 2000 g for 15 min (discarding the supernatant and ∼1 mL erythrocytes each time) to isolate purified erythrocytes. The supernatant from the first centrifugation underwent a second centrifugation at 200 g for 10 min to harvest purified PRP samples. This PRP supernatant was centrifuged sequentially at 200 g for 10 min (pellet discarded), followed by two centrifugations at 2000 g for 15 min, with the upper layer collected as PPP samples. Platelet pellets from the 2000 g spins were washed twice with 4 mL of EDTA-PBS and centrifuged at 2000 g for 15 min, then resuspended in PBS buffer. Purified PRP, PPP, erythrocytes, and platelets were quantified using an automated hematology analyzer (Shenzhen Mandray Biomedical Electronics Co., Ltd., China).

### Proteome sample preparation

For plasma samples without protein enrichment (neat), 2 μL of plasma is processed using a standard in-solution digestion protocol^2^. All chemical reagents, unless otherwise specified, were obtained from Sigma-Aldrich (St. Louis, MO, USA). Specifically, 50 μL of lysis buffer containing 8 M urea and 2 M thiourea, dissolved in 100 mM ammonium bicarbonate (ABB), is added for protein denaturation. The mixture is then treated with a final concentration of 10 mM Tris(2-carboxyethyl)phosphine (TCEP) and 40 mM iodoacetamide (IAA) in the dark for 30 min to facilitate reduction and alkylation. Subsequently, a 100 mM ABB dilution solution is used to reduce the urea concentration to below 1.2 M, followed by the addition of 1 μg of trypsin at a 1:50 ratio for overnight digestion. Finally, 30 μL of 10% Trifluoroacetic acid (TFA) is added to terminate the digestion. The resulting tryptic peptides are purified using a SOLAμ HRP 2mg/1ml 96 well plate (Thermo Fisher Scientific, Germany) and dried under vacuum. The OmniProt-based sample preparation began with 15 μL of plasma samples, which were first diluted by 75 μL of a 1×PBS buffer containing 0.05% 3-[(3-cholamidopropyl)dimethylammonio]-1-propanesulfonate (CHAPS, GPC Biotechnology GmbH, Germany) and 0.02% ammonia (referred to as buffer 2). Next, 0.5 mg of NP solution was added, and the mixture was incubated on a shaker at 30°C and 220 rpm for 30 min. After incubation, the soft corona was washed with a diluted 33% buffer 2, using centrifugation at 7000 g for 10 min, repeated three times. Finally, the soft corona was treated using the same protein denaturation, digestion, and desalting methods as the neat plasma samples. The Top14-based depletion (Thermo Fisher Scientific, Germany) method was performed as previously described^15^. Briefly, 10 μL of plasma samples were depleted of 14 high-abundance proteins using human affinity depletion resin and then concentrated into 50 μL through a 3 kDa MWCO Ultra Centrifugal Filter (Merck, Germany). Next, remaining protein was denatured in 8 M urea at 30°C for 30 min, reduced with 10 mM TCEP, and alkylated with 40 mM IAA. Subsequently, 75 µL of 100 mM ABB solution was added to dilute the urea concentration. Trypsin was introduced at 1.0 µg to initiate proteolytic digestion. The resulting tryptic peptides were desalted via SOLAμ HRP 2mg/1ml 96 well plate (Thermo Fisher Scientific, Germany) and stored at −80°C for downstream analyses.

### Enrichment of plasma proteins using various NPs for building a matching library

We utilized pooled plasma samples from patients with lung cancer to enrich low-abundance proteins using various NPs, following the OmniProt method. Approximately 200 µg of plasma digests were loaded onto a Waters XBridge column (2.1 mm × 150 mm, BEH C18, 5 μm) with a Thermo Scientific UltiMate™ 3000 RSLC LC system equipped for basic-pH reverse phase liquid chromatography (basic pH RPLC), producing 120 fractions that were subsequently combined into either 30 or 60 fractions. These were evaporated and resuspended for further analysis. Each fraction was loaded into a trap column and separated on a custom-made analytical column (75 μm i.d. × 15 cm, 1.9 μm) using a 24-min LC-MS method with a Vanquish™ Neo UHPLC system (Thermo Fisher Scientific, Germany). The initial gradient began at 8% buffer B (buffer B: 80% MS-grade ACN with 0.1% (v/v) formic acid; buffer A: 98% MS-grade ACN with 0.1% (v/v) formic acid), increasing to 10% B over 1.5 min, then to 30% B over 16 min, and finally to 40% B over 2 min, with 4.3 min reserved for column cleaning and equilibration. The eluted peptides were analyzed using an Orbitrap Astral mass spectrometer (Thermo Fisher Scientific, Germany), resulting in 750 DIA files for spectral library construction.

### DIA acquisition

First, approximately 300 ng peptide was loaded into a trap column and subsequently separated on an analytical column (75 μm i.d. × 15 cm, 1.9 μm, custom-made) using a 24-min LC-MS method. The initial liquid phase gradient for LC separation commenced with 8% buffer B (buffer B: 80% ACN with 0.1% (v/v) formic acid; buffer A: 98% MS-grade ACN with 0.1% (v/v) formic acid), increasing to 10% B over 1.5 min, then to 30% B over 16 min, and finally to 40% B over 2 min. The LC method incorporated a 4.3-min column cleaning and equilibration phase prior to subsequent runs. The eluted peptides were analyzed using an Orbitrap Astral mass spectrometer (Thermo Fisher Scientific, Germany), with FAIMS voltage set at −42 V, a full scan resolution of 240,000, and a scan range of 380–980 Th. MS/MS scans were conducted within the same range, employing a DIA isolation window of 2 Da. The mass spectrometry parameters for library construction were largely consistent with those used for routine DIA acquisition, except that the isolation window was set to 1 Da.

### DIA-MS analysis

DIA files were analyzed with DIA-NN^22^ (v1.8.1). For platelet-, erythrocyte-, and coagulation-related DIA files, analyses were conducted against the reviewed UniProt Human database (20,377 entries, downloaded in May 2020) with default parameters and Match-Between-Runs (MBR) enabled. For all other DIA datasets, a custom spectral library comprising 126,661 peptide precursors mapped to 10,109 protein groups (generated in this study) was employed, using identical default settings and MBR. A 1% false discovery rate (FDR) threshold was applied at both peptide precursor and protein levels, with quantification performed in Robust LC mode to ensure analytical precision. DIA files generated from the human and bovine plasma benchmarking experiment were analyzed using DIA-NN (v1.8.1) against a combined database comprising the reviewed UniProt Human database (20,377 entries, downloaded in May 2020) and the reviewed UniProt Bovine database (6048 entries, downloaded in January 2025), with default parameters and MBR.

### Bioinformatics analysis

The statistical analysis of the data was performed using R software (v4.1.2) and Python (v3.12.5 and v3.10), which included the use of heatmap and R package plot functions. The proteins in the heatmaps were hierarchically clustered using the centroid method. The main report containing peptide and protein quantification results was retained for downstream evaluation. OmniProt workflow precision was assessed by calculating the CVs for each protein across biological replicates. We used bovine-specific peptide precursors to evaluate the accuracy of the OmniProt workflow. Peptide precursor intensities were log-transformed, and technical variations across replicates were corrected through global median normalization of log-intensities. Then we compared measured bovine-specific peptide precursors’ fold changes of spiked-in ratios with their expected theoretical values.

To identify platelet- and erythrocyte-related biomarkers, proteins with missing values in more than 50% of samples were first excluded. Candidate markers for platelet or erythrocyte contamination were selected from the top 100 proteins most strongly correlated with platelet concentration changes. Among these, the 30 proteins showing the highest protein abundance were prioritized as potential biomarkers. Candidate markers for coagulation-related contamination were identified through the following workflow. First, proteins with missing values in more than 50% of samples were removed. The abundance of each protein under each condition was calculated as the mean of three biological replicates. Next, proteins meeting adjusted p-value < 0.05 and fold change > 10 between plasma and serum samples were selected. From this subset, the top 20 most abundant proteins were prioritized as potential coagulant contamination markers.

### Development of Baize software

The Baize algorithm operates through a unified computational workflow designed for automated contamination assessment. Users input a protein matrix (generated by search engines such as DIA-NN), and the software calculates contamination index for platelets, erythrocytes, and coagulation-related protein using predefined biomarker panels (e.g., PF4 for platelets, HBA1 for erythrocytes, FGA for coagulation). These indexes are derived by normalizing the summed intensities of contamination-specific biomarkers against the total plasma proteome intensity. The output provides each sample’s contamination profiles, enabling rapid quality control decisions without requiring manual intervention.

## Data availability

OmniProt is developed in Python and is freely available. The latest software version is available at https://www.guomics.com/Baize. The mass spectrometry raw data have been deposited with the ProteomeXchange Consortium via the PRIDE^56^ partner. The dataset will be made publicly available upon acceptance of the manuscript.

## Code availability

Code relevant for data analysis in this study is available at GitHub [https://github.com/guomics-lab/Baize].

## Acknowledgments

This work is supported by grants from the “Pioneer” and “Leading Goose” R&D Program of Zhejiang (2023C03056), the National Natural Science Foundation of China (Key Joint Research Program, Grant No. U24A20476), National Key R&D Program of China (Grant No. 2022YFF0608403 to Dr Yi Zhu), the “Pioneer” and “Leading Goose” R&D Program of Zhejiang (2024SSYS0035). Fig.1a, Fig. 2a, Fig. 4a, Fig. 4f, Fig. 4i, Fig. 5a and Fig. 5f were created with Biorender.com. The language of the first draft has been polished with DeepSeek R1 before proofreading by the co-authors.

## Author contributions

T.G., and H.G. conceived the project and wrote the manuscript. Y. Zhan, H.G., Y. Zheng, Q.L., and D.X. prepared the samples, Z. Zhu, W.G., Z. Zhang, and H.G. participated in data analysis. Z.X., Y.Z., and H.G. developed the Baize software. Y.L., H.G., and L.Z. collected the blood samples from the hospital. J.L., and J.Z., provided the Zeolite NaY nanoparticle. R.S. provided the hematology analyzer. H.M., and Y. Zhu. revised this manuscript.

---

**Supplementary Figure 1.**
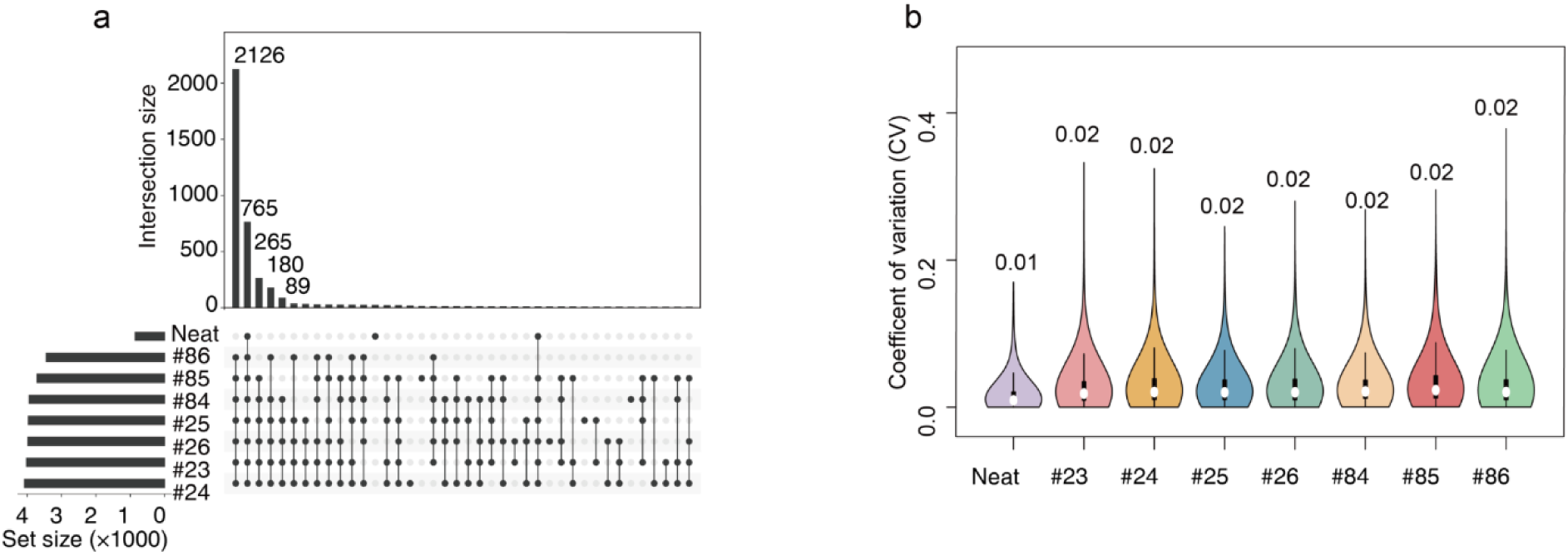
Comparative analysis of various SiO_2_-based nanoparticles. (a) Upset diagram of identified protein groups across various SiO_2_-based NPs. (b) Coefficients of variation for identified protein groups across various SiO_2_-based NPs.

**Supplementary Figure 2.**
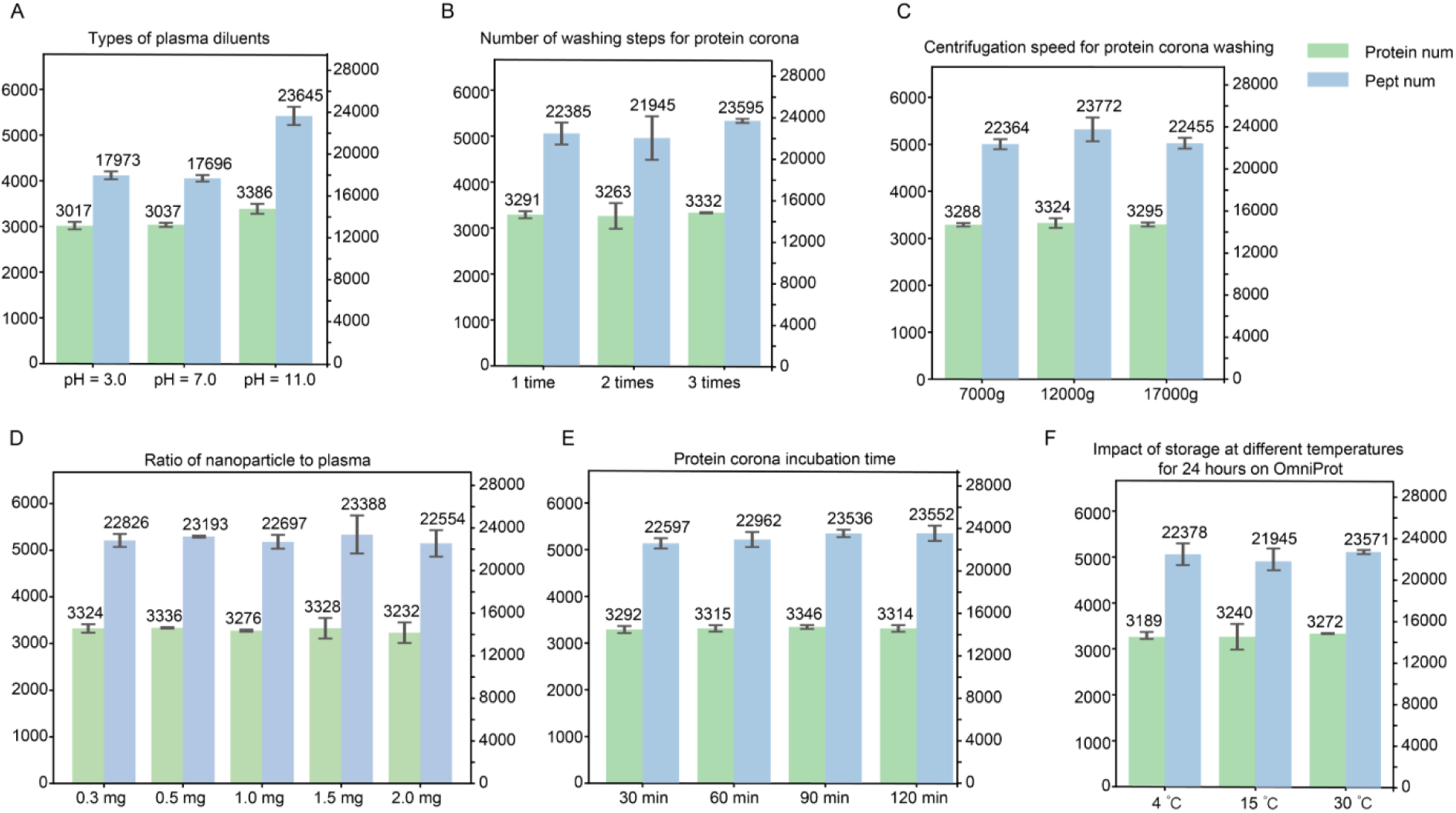
The optimization of parameters influencing protein corona formation. The parameters influencing protein corona formation included type of plasma diluents (A), number of washing steps for protein corona (B), centrifugation speed for protein corona washing (C), ratio of nanoparticle to plasma (D), protein corona incubation time (E), and impact of storage at different temperatures for 24 hours on OmniProt.

**Supplementary Figure 3.**
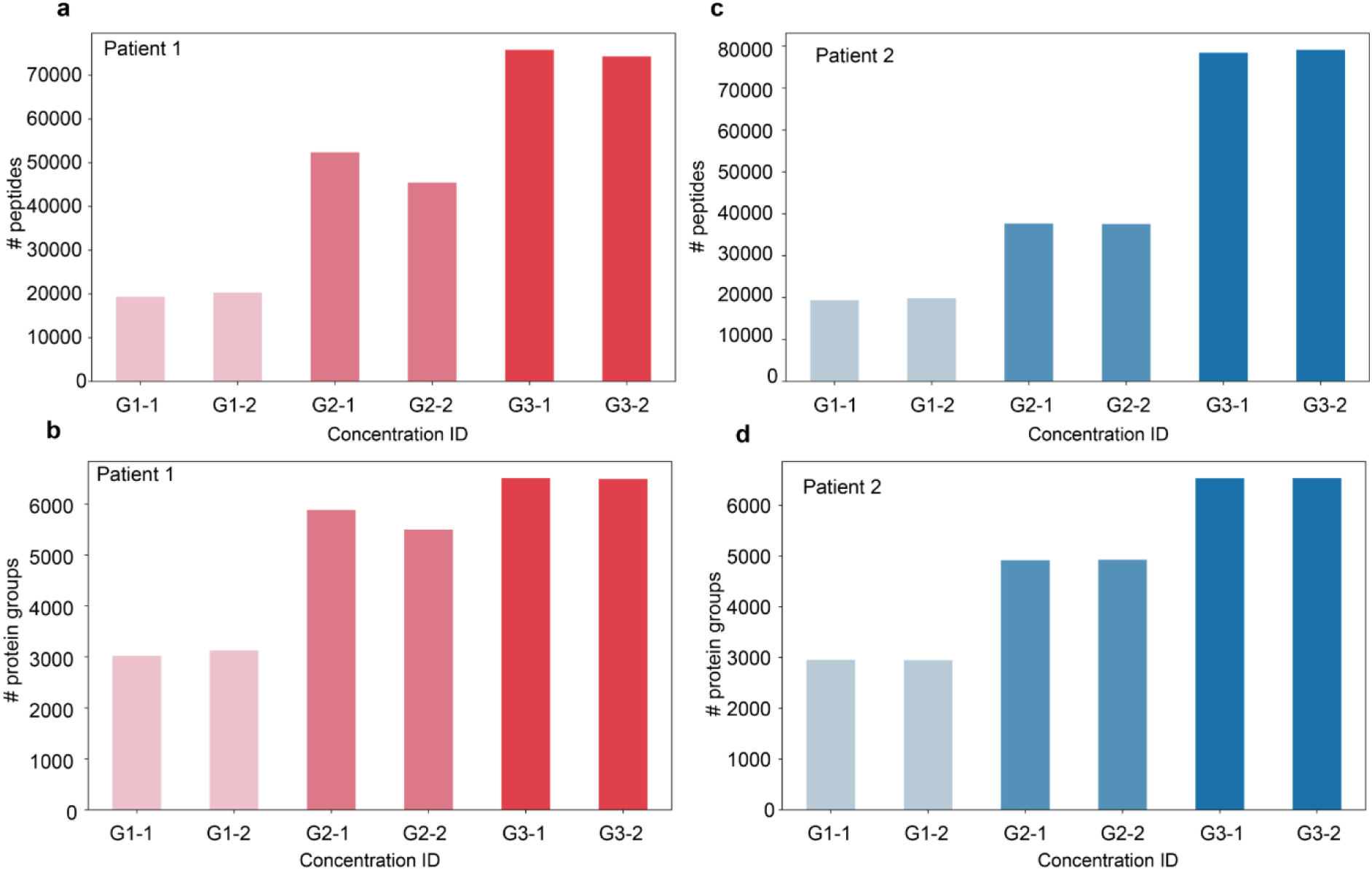
The identified peptides and protein groups in various plasma samples. Number of identified peptides (a) and protein groups (b) in plasma samples from patient 1. Number of identified peptides (c) and protein groups (d) in plasma samples from patient 2.

**Supplementary Figure 4.**
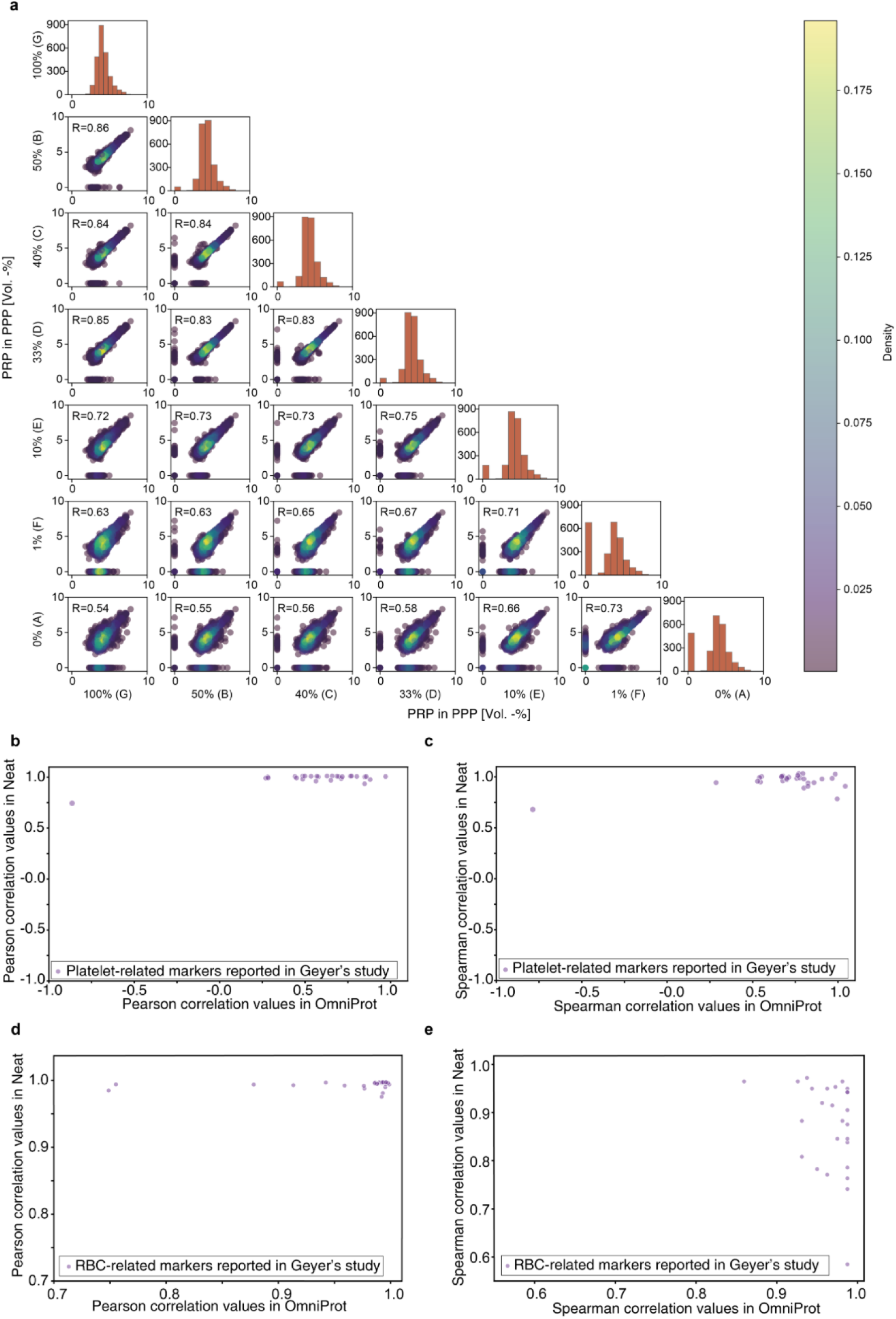
Correlation of platelet/erythrocyte-related markers from Geyer’s study in neat and OmniProt plasma samples. (a) Comparative Pearson correlation analysis of plasma proteome across varied PRP-to-PPP ratios. Pearson (b) and Spearman (c) correlation of platelet-related markers between neat and OmniProt samples. Pearson (d) and Spearman (e) correlation of RBC-related markers between neat and OmniProt samples.

**Supplementary Figure 5.**
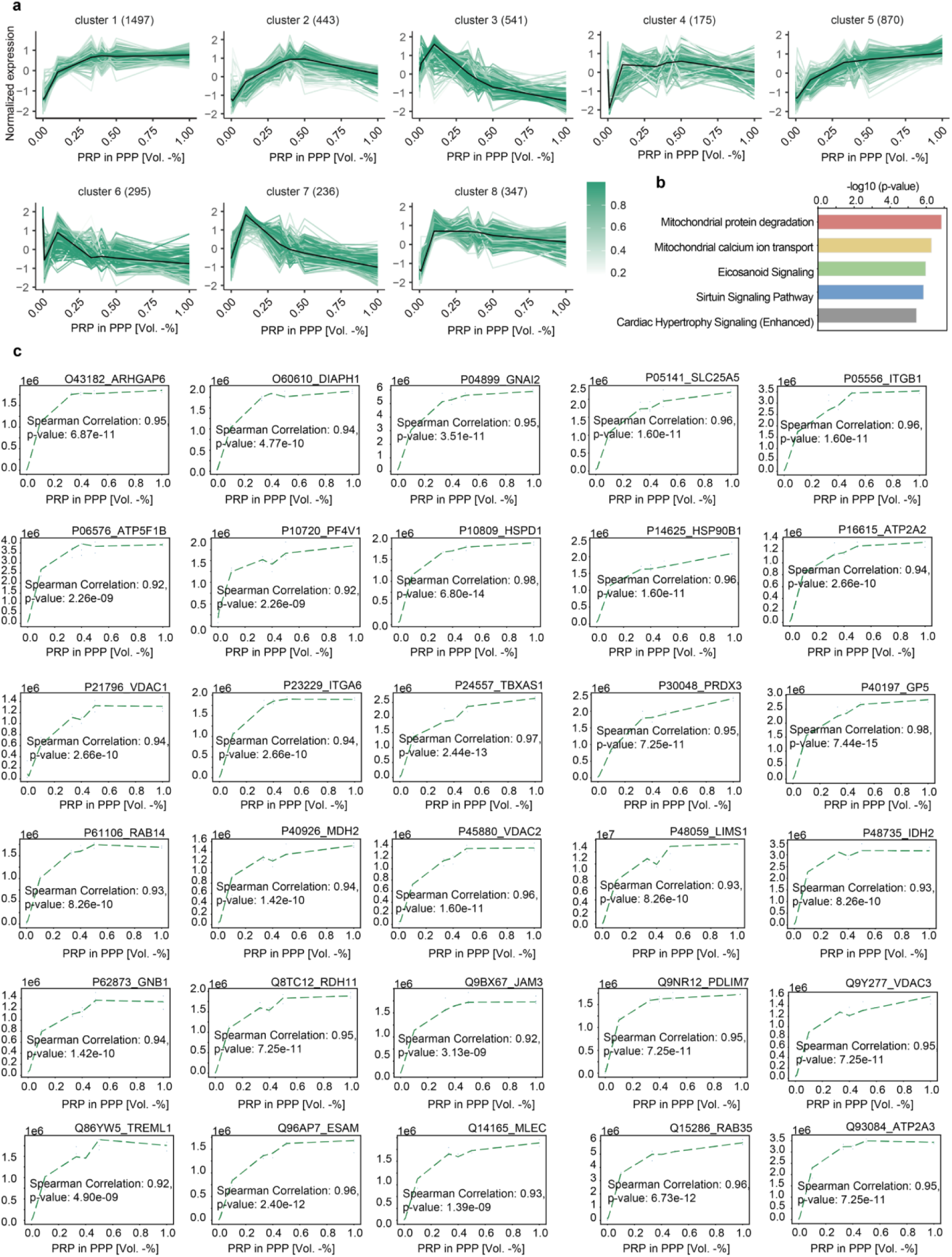
Correlation of platelet-related markers from Geyer’s study in plasma samples prepared using the neat and OmniProt workflows. (a) Mfuzz analysis of identified proteins with missing rate <50%. (b) Spearman correlation for 30 markers in discovery dataset.

**Supplementary Figure 6.**
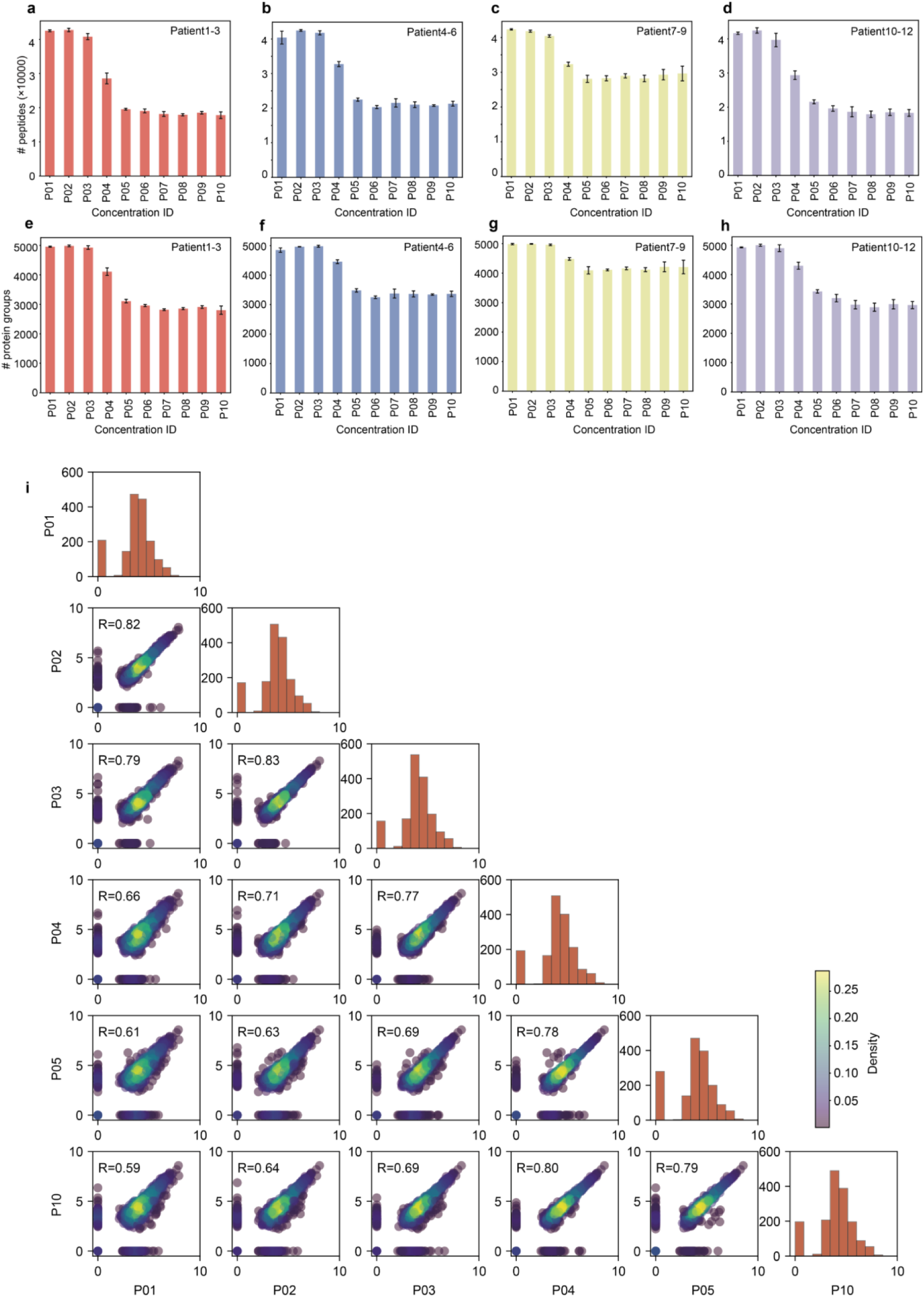
Number of identified peptide precursors (a-d) and protein groups (e-h) across four patients.

**Supplementary Figure 7.**
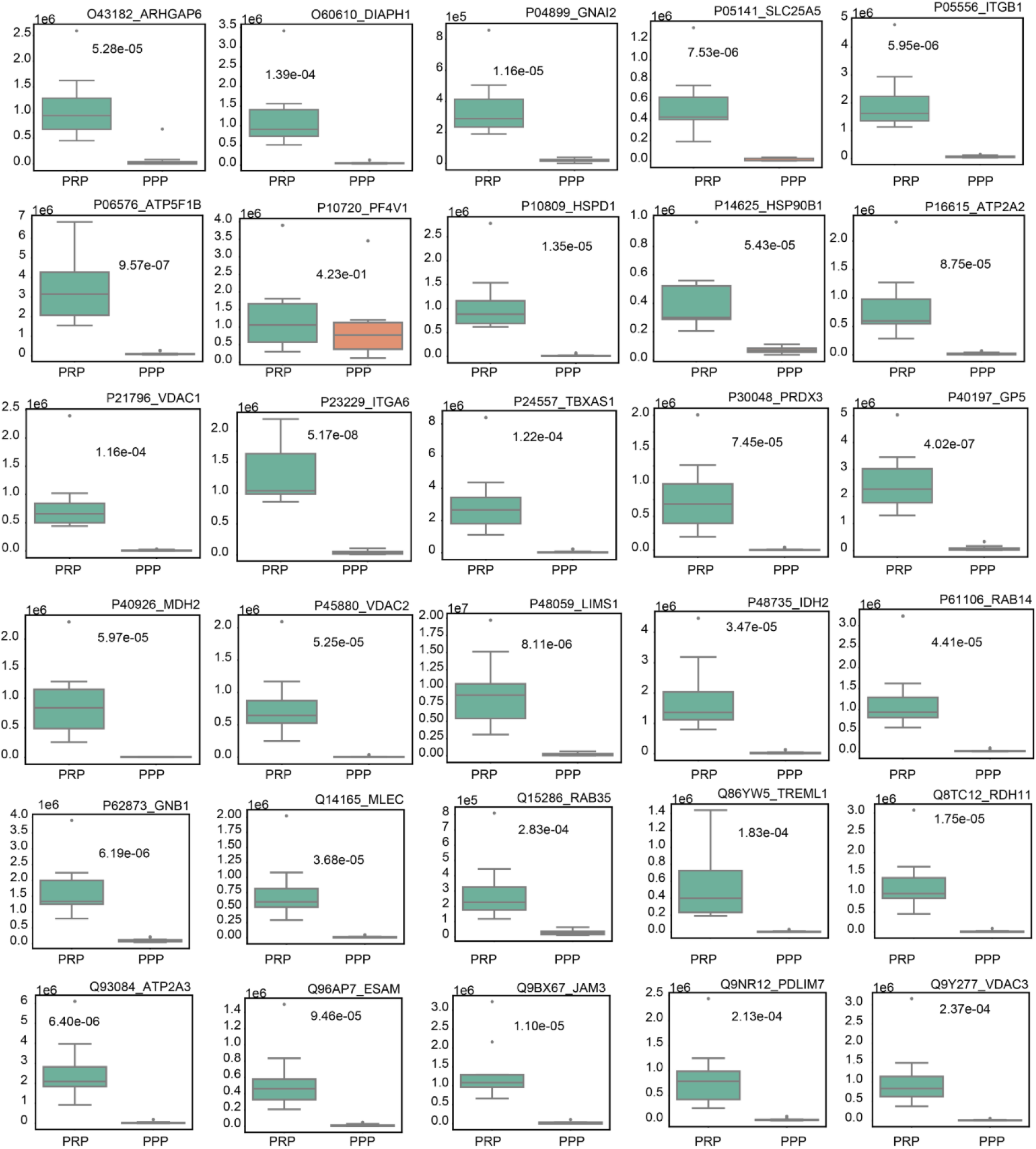
Comparative analysis of protein abundance distributions in matched PRP and PPP samples. The p values were assessed by paired Student’s t-test for comparisons.

**Supplementary Figure 8.**
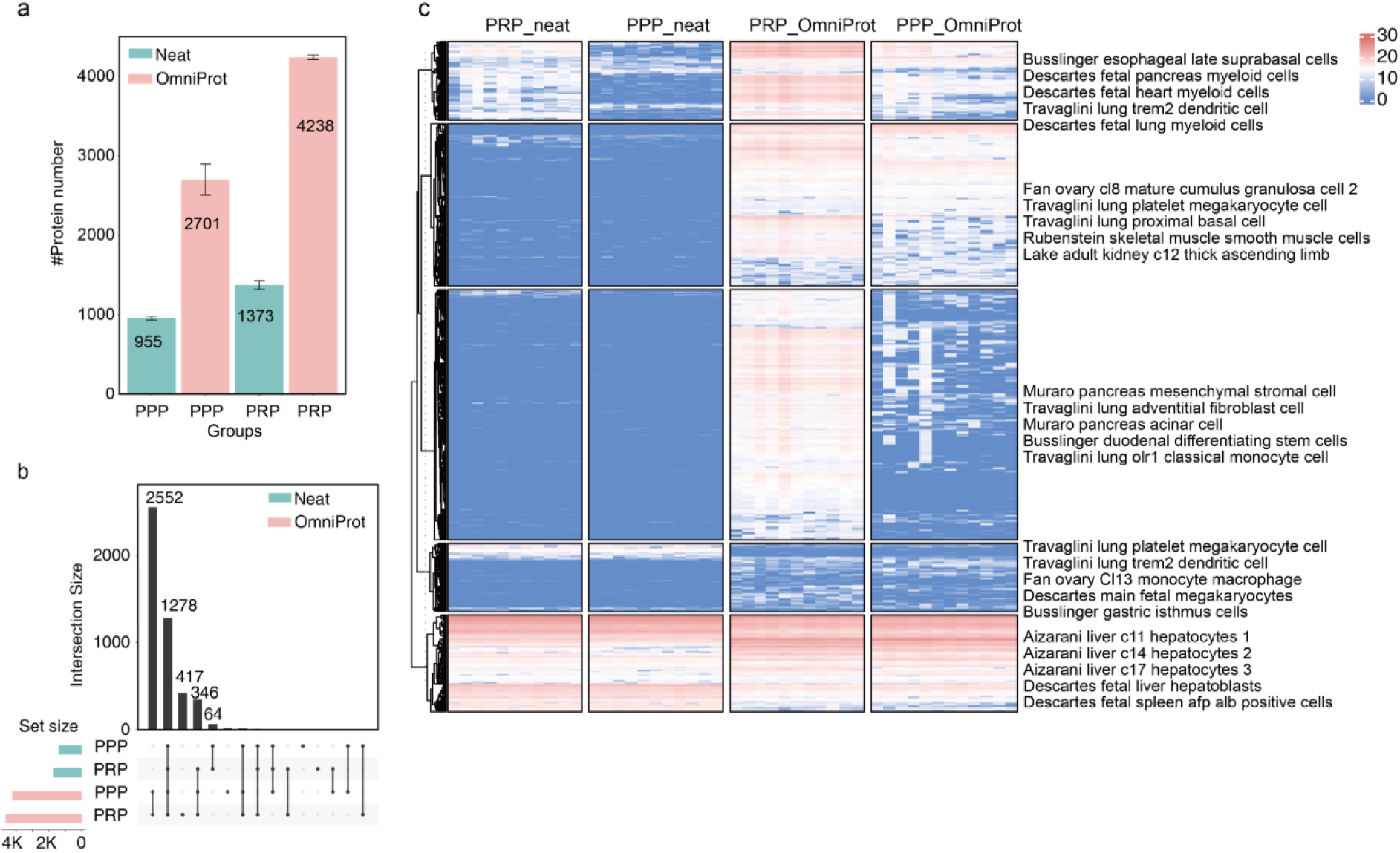
Heatmap of protein identification in 11 paired PRP and PPP samples. (a) Number of proteins identified in 11 PRP and PPP patient samples under Neat and OmniProt methods. (b) Upset Plot comparing protein Identification distribution in 11 PRP and PPP patient samples under Neat and OmniProt methods. (c) Heatmap of protein identification in 11 paired PPP and PRP samples.

**Supplementary Figure 9.**
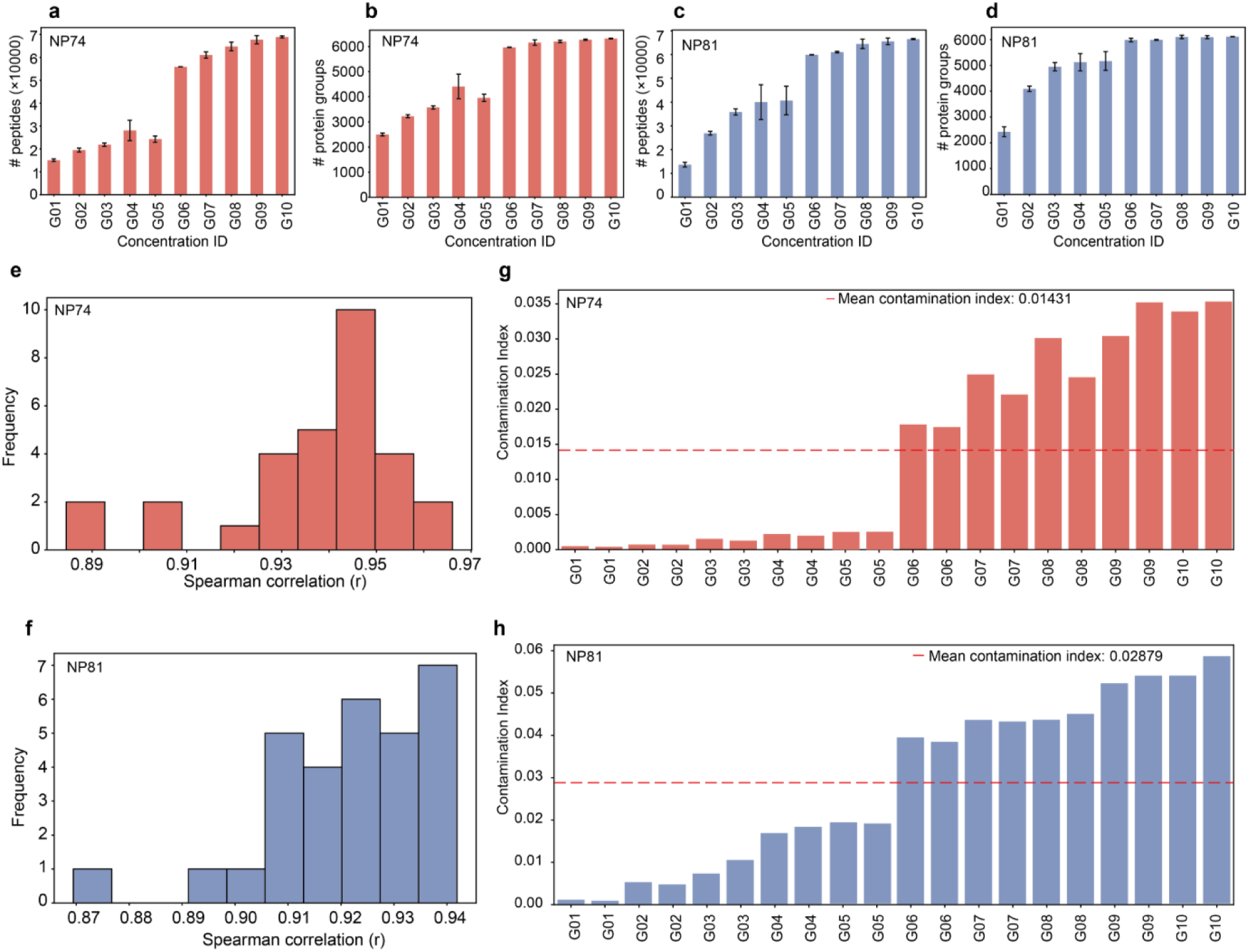
Performance of nanoparticle-based methods in processing plasma samples with varying platelet contamination levels. Number of identified peptide precursors (a) and protein groups (b) in NP74. Number of identified peptide precursors (c) and protein groups (d) in NP81. Spearman correlation of 30 platelet-related biomarkers in NP74-processed samples (e) and NP81-processed samples (f) across platelet contamination levels. Contamination index profiles of NP74-processed samples (g) and NP81-processed samples (h) with graded platelet contamination.

**Supplementary Figure 10.**
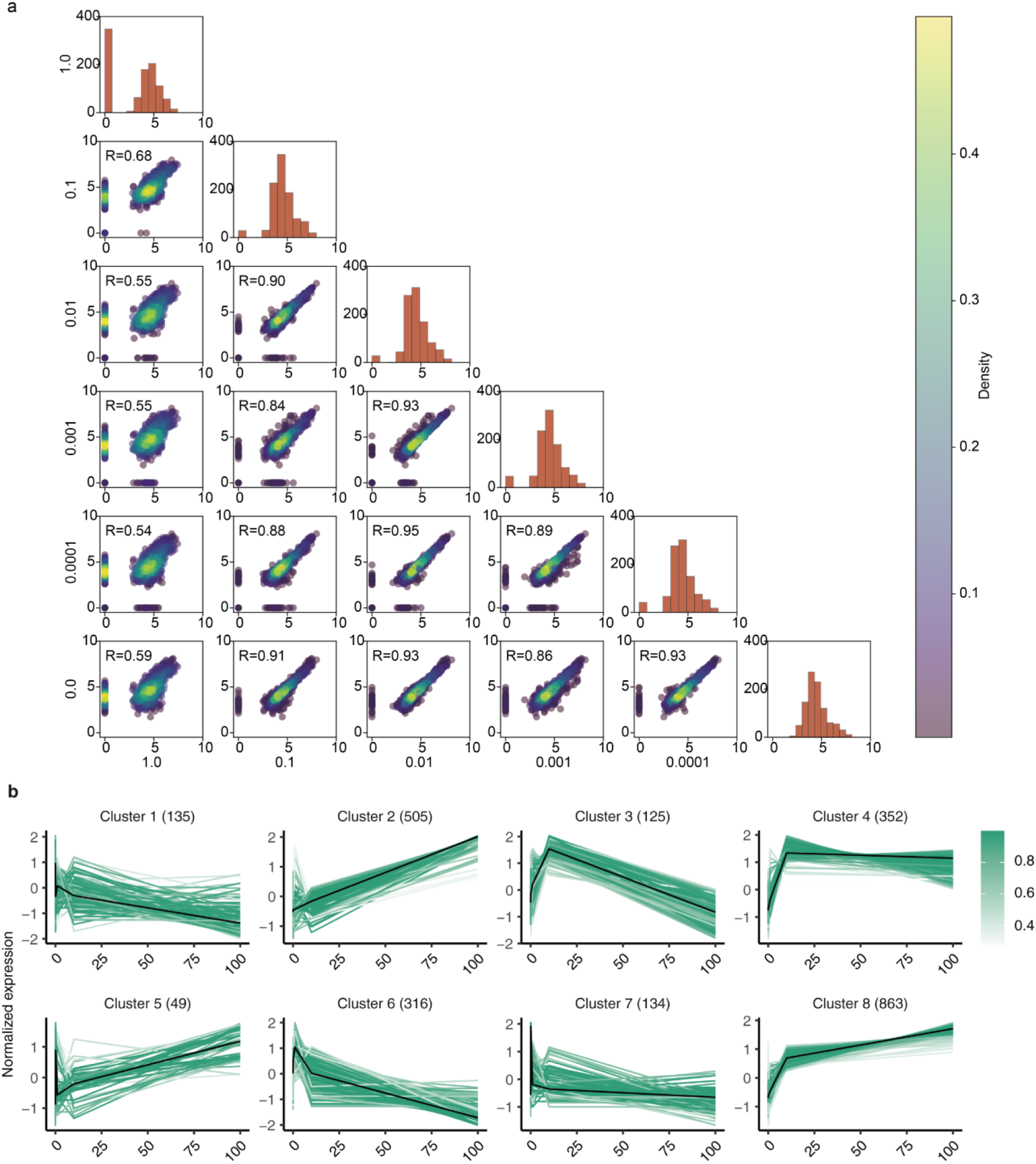
Performance of erythrocyte-derived protein contamination in OmniProt plasma samples. (a) Pearson analysis of plasma proteome integrity across erythrocyte contamination gradients. (b) Mfuzz analysis of identified proteins with missing rate <50%.

**Supplementary Figure 11.**
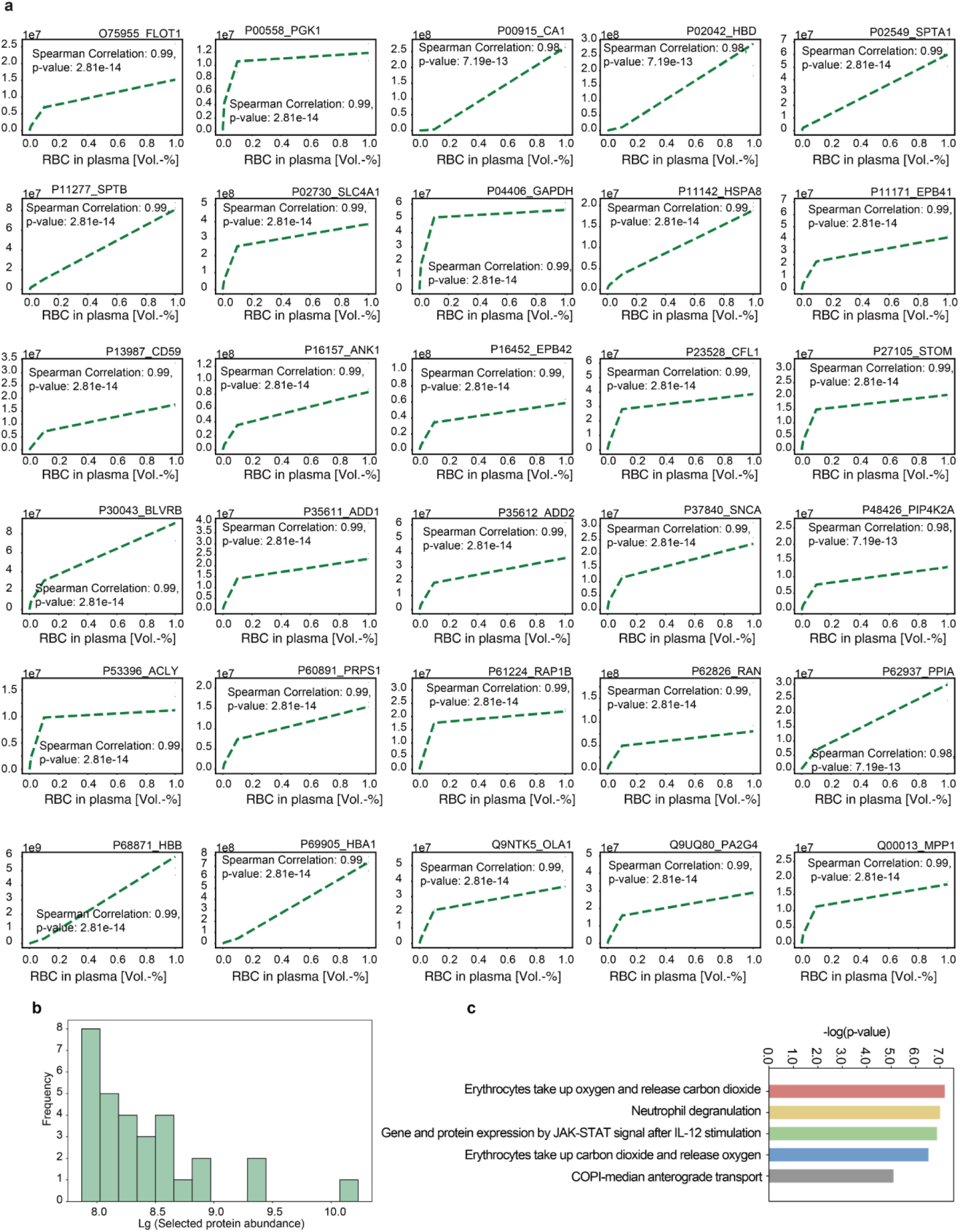
Correlation of erythrocyte-related markers in OmniProt plasma samples. (a) Distribution of protein abundance across selected 30 platelet-related markers. (b) Abundance distribution of erythrocyte-associated markers in this study. (c) Comparative abundance of erythrocyte markers from prior study.

**Supplementary Figure 12.**
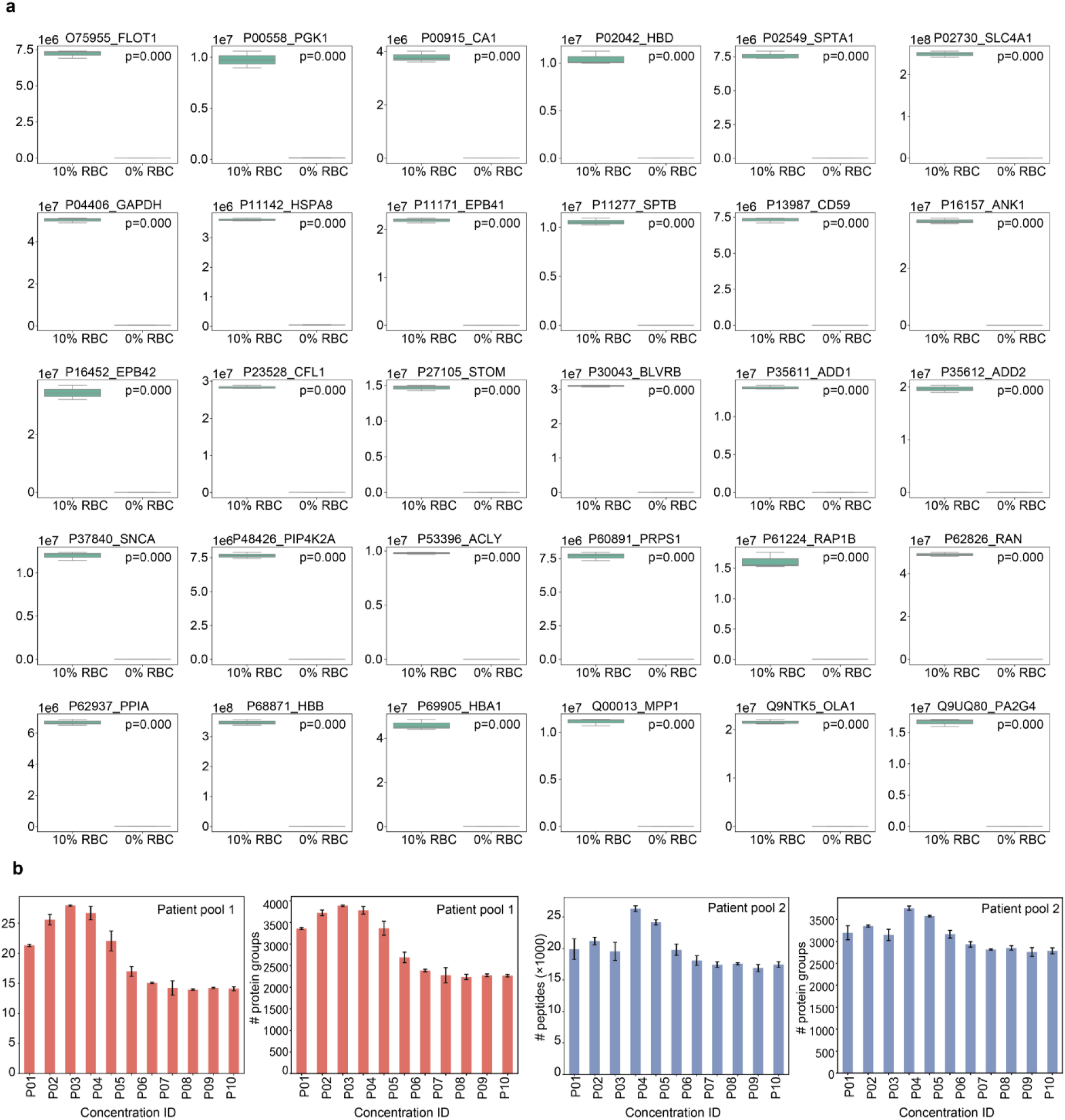
Performance evaluation of 30 RBC-related markers in the discovery set. (a) Comparative analysis of protein abundance distributions in 10% RBC and 0% RBC samples (b) Number of identified peptide precursors and protein groups across two patient pools. The p values were assessed by paired Student’s t-test for comparisons.

